# A screen for stress-induced sleep genes in *C. elegans* reveals a role for glutamate signaling

**DOI:** 10.64898/2026.03.02.709123

**Authors:** Quinn Howe, Caroline Kominick, Mercedes Pierce, Gina Gazarra, Caroline E. Curtin, Santino Diana, Fredy Abboud, Jordan Olenginski, Mary Frattara, Ting Brown, Teagan McCarthy, Paige Conrad, Lauren Yoslov, Rachel Vemula, Anthony Gargani, Emily Le, Matthew D. Nelson

## Abstract

Sleep is an essential behavioral state that evolved early in animals, possibly with the advent of the nervous system. The complexity of sleep neural networks varies significantly across phylogeny, yet common signaling molecules exist. Stress-induced sleep (SIS) of *Caenorhabditis elegans* is controlled by two sleep interneurons (ALA and RIS), within a 302-celled nervous system. Even in this simple framework, a complex array of signaling molecules is expressed. Here, we surveyed some of these genes for roles in SIS. These included neuropeptides, g-protein coupled receptors, a two-pored potassium channel, and glutamate signaling components. We found that multiple genes are required for sleep maintenance (i.e., amounts), and/or the precise timing of sleep initiation. In particular, we identified an important role for glutamate signaling. The conserved ionotropic glutamate receptor *glr-5*, regulates sleep maintenance and timing, and is required in a 3-celled circuit of interneurons connected by gap junctions and chemical synapses with RIS. This work suggests that numerous redundant and/or parallel mechanisms have evolved to modulate a simple sleep-regulating circuit in *C. elegans*, and we speculate that conserved pathways may play similar roles in animals with more complex systems.

## Introduction

Sleep exists in all animals with nervous systems (Anafi *et al*. 2019), and may even have evolved in ancestral animals who lacked true tissues, considering parazoans express circadian and synaptic transmission genes (Rivera *et al*. 2012; Musser *et al*. 2021). Conserved sleep regulatory molecules are found even in primitive animals with radially-arranged nerve nets, like hydra (Kanaya *et al*. 2020) and jellyfish (Nath *et al*. 2017), suggesting that this mysterious behavioral state evolved once, prior to the appearance of the bilaterians. Thus, conserved mechanisms may regulate sleep across phylogeny (Trojanowski and Raizen 2016; Anafi *et al*. 2019), despite dramatic differences in nervous system complexity. Invertebrate models have emerged as powerful tools for sleep gene discovery, and mechanistic dissection.

The nematode *Caenorhabditis elegans* has a 302 cell, fully-mapped nervous system (White *et al*. 1986; Taylor *et al*. 2021), and displays sleep controlled by conserved genes (Honer and Nelson 2024). Stress-induced sleep (SIS) is a particularly intriguing model for sleep gene identification, considering it’s experiment reproducibility, and control by just two interneurons, ALA (Hill *et al*. 2014) and RIS (Konietzka *et al*. 2020). SIS provides restorative functions by allowing for reallocation of energetics from excitable cells towards cellular repair, akin to sickness sleep of mammals (Davis and Raizen 2017). Extreme temperatures, ethanol, pore-forming toxins, hyperosmotic conditions (Hill *et al*. 2014), wounding (Goetting *et al*. 2020), exposure to the bacterial toxin indole (Diya *et al*. 2025), viral infection (Iannacone *et al*. 2024), and ultraviolet irradiation (Debardeleben *et al*. 2017), can lead to SIS. Noxious stimuli can damage tissues differently, but a common behavioral program occurs: 1) avoidance and escape; 2) sleep; 3) arousal. The timing and amount (i.e., maintenance) of sleep depends on the extent of cellular damage (Hill *et al*. 2014), thus temporal dynamics can vary.

Numerous signaling mechanisms must coordinate the opposing states of sleep and avoidance. Some of the most critical molecules of sleep regulation are epidermal growth factors coded by *siss-1* (Hill *et al*. 2024), which are shed from damaged tissues to activate EGF-receptors, coded by *let-23*, on the surface of the ALA (Van Buskirk and Sternberg 2007), and RIS (Konietzka *et al*. 2020). Wounding of skin promotes ALA and RIS activation by *siss-1* as well as by the upregulation and release of antimicrobial peptides (Pujol *et al*. 2008; Sinner *et al*. 2021). ALA and RIS activation leads to the secretion of a collection of neuropeptides required for the sleep behaviors. These include *nlp-8*, *nlp-14*, *flp-13*, and *flp-24*, expressed in the ALA (Nelson *et al*. 2014; Nath *et al*. 2016; Honer *et al*. 2020), and *flp-11* in the RIS (Turek *et al*. 2016; Konietzka *et al*. 2020). Upon secretion, these peptides may function to inhibit wake and motor interneurons (Iannacone *et al*. 2017; Cianciulli *et al*. 2019), and cholinergic motor neurons (Rossi *et al*. 2025), via inhibitory G-protein coupled receptors (GPCRs) (Trojanowski *et al*. 2015; Iannacone *et al*. 2017). Despite the identification of neuropeptides, only a small subset of GPCRs have been described. The GPCR *dmsr-1* functions as a *flp-11* receptor on the RIS, which provides an autocrine feedback mechanism to regulate sleep duration, as well as a *flp-11* receptor on excitatory cholinergic neurons (Rossi *et al*. 2025), and a *flp-13* receptor on wake interneurons (Iannacone *et al*. 2017). The orphaned GPCR *npr-38* is required for sleep in the ADL sensory neurons, and may play a broader timing role for SIS. Overexpression (OE) of *npr-38* causes a profound shift in the timing of sleep initiation, in that it occurs early when arousal and escape should dominate. Additionally, *npr-38* is required for heat and blue light avoidance behaviors, as well as general arousal (Le *et al*. 2023). Thus, *npr-38* may regulate the timing of ALA and RIS activation, together with other unknown pathways, to coordinate the broader stress response.

Only a small number of other cells have been implicated in SIS, which include ADL, AIY, RMG, DVA, and RIF, as well as cholinergic neurons. The AIY is one site of action for *dmsr-1* (Iannacone *et al*. 2017), and *dmsr-1* also functions on RIS, and cholinergic neurons (Rossi *et al*. 2025). RMG promotes arousal from SIS, possibly via the secretion of wake-promoting *pdf-1* neuropeptides (Choi *et al*. 2013; Soto *et al*. 2019). Mutants of the neuropeptide receptor *npr-1* display very little SIS under certain conditions (Soto *et al*. 2019); *npr-1* functions in the RMG (Choi *et al*. 2013). Thus, it appears that the RMG can override the action of the ALA and RIS to promote arousal. The DVA and RIF also stimulate arousal; optogenetic activation of the cAMP pathway in either of these cells induces waking, and impairs sleep. Also, cAMP levels are reduced in DVA and RIF during sleep (Cianciulli *et al*. 2019). Thus, the ADL, AIY, RMG, DVA, and RIF neurons modulate sleep in different ways, how they communicate with that ALA and RIS mechanistically is still unclear.

Many questions surrounding SIS still remain fully answered. First, how do animals coordinate the temporal aspects of stress avoidance and escape, with the opposing events of sleep neuron activation? This is an essential balance, considering animals must remove themselves from dangerous environments before entering a recovery sleep state which will render them vulnerable to the initial threats. Next, how do sleep neurons mechanistically induce the behavioral aspects of SIS? Data suggest that the activation of GPCRs leads to the mobilization of inhibitory G-proteins, and thus the modulation of second messenger system, however, little mechanistic data has been shown to support such a model. The identification of new receptors and signaling molecules will be an important first step in expanding our mechanistic insight into these events. Last, what signals promote arousal from SIS, and how do they over-ride the effects of sleep neuron activation?

As a first step to better understanding these mechanisms, we sought to identify new sleep regulating molecules by measuring sleep in mutants of genes expressed in the abovementioned cells, as well as a few others connected to them by chemical synapses or gap junctions (**Figure 1**) (White *et al*. 1986). These included neuropeptides, GPCRs, a potassium channel, and glutamate signaling molecules (**Figure 1 and Table S1**). Loss of function of some of the genes reduced sleep, while others enhanced it, and in some cases we observed an alteration in the timing of sleep. We identified a particularly important role for the glutamate receptor *glr-5*, in which mutants display significantly less sleep and initiated sleep later than controls. Using cell-specific rescue we propose a model in which *glr-5* functions in the RIS to modulate sleep timing, and in the RIS, AIB, and RIM interneurons to regulate sleep amounts (i.e., sleep maintenance).

**Figure 1:**
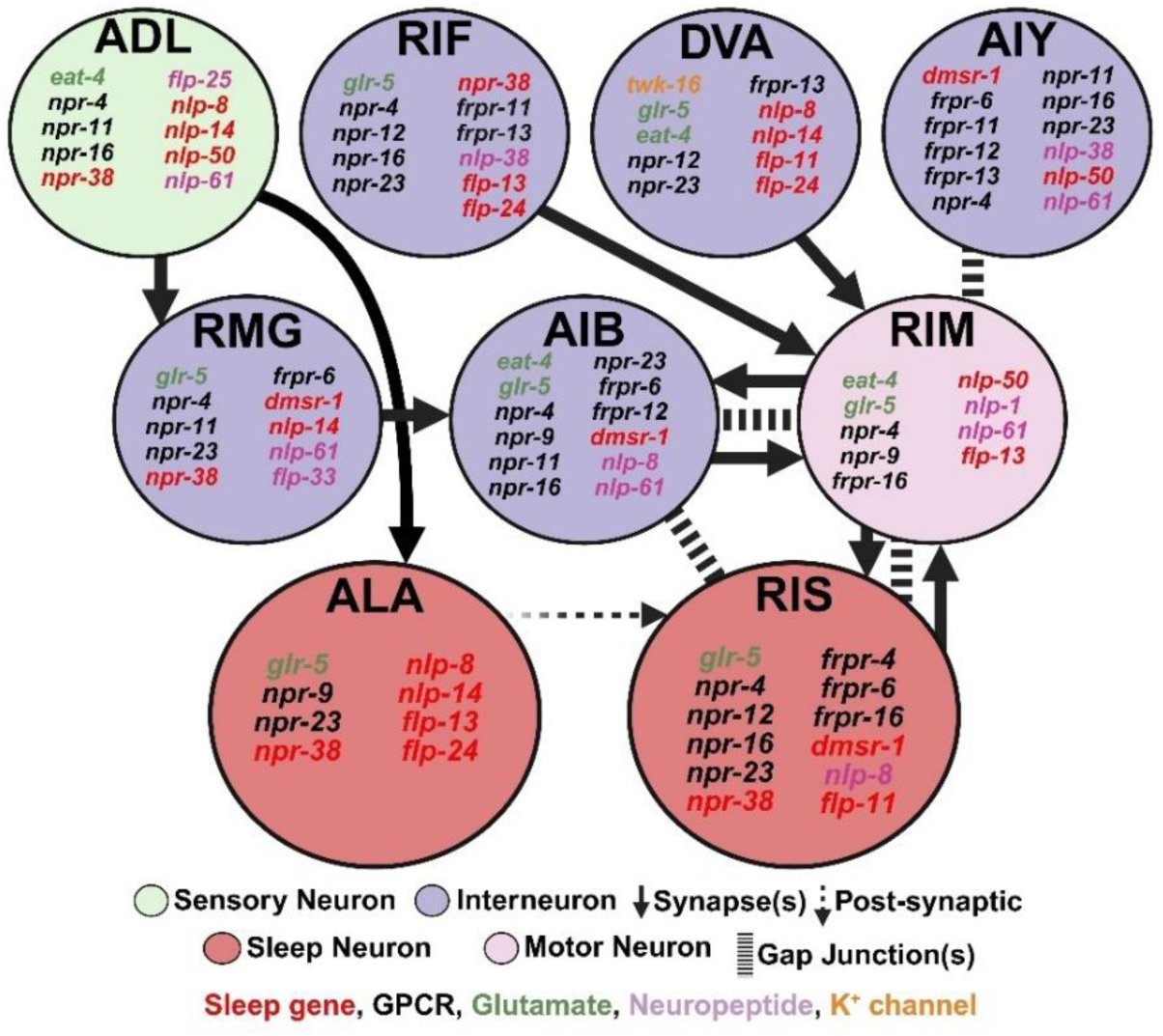
Expression of signaling molecule genes in a sleep-regulating circuit. Expression is based on published RNAseq data (Taylor *et al*. 2021), chemical synapses are indicated by solid arrows, gap junctions by dashed lines (White *et al*. 1986), and post-synaptic communication by a dashed arrow (Konietzka *et al*. 2020).

## Results

### Multiple neuropeptide genes are required for stress-induced sleep

Numerous neuropeptide genes are expressed in this simple circuit (**Figure 1**), we focused on a subset of them, for a few reasons. First, *flp-3*, *flp-25*, and *flp-33* code RFamide peptides like *flp-11* and *flp-13* (Nathoo *et al*. 2001); RFamides play conserved sleep roles across phylogeny (Lenz *et al*. 2015; Nath *et al*. 2016; Turek *et al*. 2016; Lee *et al*. 2017). We focused on the genes *nlp-1*, *nlp-38*, and *nlp-61* due to their high levels of expression in some of these cells (Taylor *et al*. 2021), and potential candidates as ligands for *npr-38* (Personal communication, Isabel Beets). Loss-of-function alleles were available for each gene except *nlp-61*, which we constructed using CRISPR. We focused on UV-sleep because it is highly temporally reproducible (Debardeleben *et al*. 2017), thus allows for the detection of timing defects.

We measured sleep using the WorMotel system (Churgin *et al*. 2017) in *flp-13*(*tm2427*) deletion mutants, in order to establish a timing and sleep maintenance baseline in a known sleep-defective mutant (Nelson *et al*. 2014). As expected, total sleep levels were significantly lower in *flp-13* mutants compared to the wild type, and reduced at multiple time points between 60-200 minutes post UV exposure. Additionally, the peak of sleep induction was shifted to a later point than controls (**Figure 2A**). Next, we measured UV-sleep in mutants for *flp-3* (Expressed in PQR), and *flp-33* (Expressed in RMG), but did not observe any differences from the wild type (**Figure 2B**). However, *flp-3*; *flp-33* double mutants showed subtle but significant alteration in sleep timing; the sleep program was delayed, with a significant difference in sleep at 70- and 80-minutes post-UV stress. The total amounts of sleep were unchanged in these double mutants (**Figure 2C**). These data suggest that *flp-3* and *flp-33* may function with *flp-13* to promote sleep at the early stages of sleep. Next, we tested sleep in mutants for *flp-25* (Expressed in ADL and PQG), surprisingly, *flp-25* mutants displayed a significant increase in sleep, primarily due to enhancement at 80-120 minutes post-stress (**Figure 2D**). These data indicate that *flp-25* peptides promote arousal from sleep. Two different *nlp-1* (Expressed in RIM) mutant strains were examined, and both showed no changes in total sleep, however, each displayed a reproducible shift in the timing of SIS (**Figures 2E and F**). This early sleep phenotype was reminiscent of sleep in *nlp-14* mutants (Honer *et al*. 2020), and following *npr-38* OE (Le *et al*. 2023). Thus, *nlp-1* may modulate sleep timing. Both *nlp-38* (Expressed in AIY and RIF) and *nlp-61* (Expressed in ADL, AIB, AIY, RIM, and RMG) mutants displayed reductions in sleep, however in different ways. Loss of *nlp-38* caused a small reduction in sleep at 80- and 100-minutes post-stress but not in total levels (**Figure 2G**). *nlp-61* mutants displayed a significant reduction in total sleep, with levels lower at 60-, 80-, and 90-minutes post-stress (**Figure 2H**). Thus, *nlp-38* and *nlp-61* may play small sleep-promoting roles.

**Figure 2:**
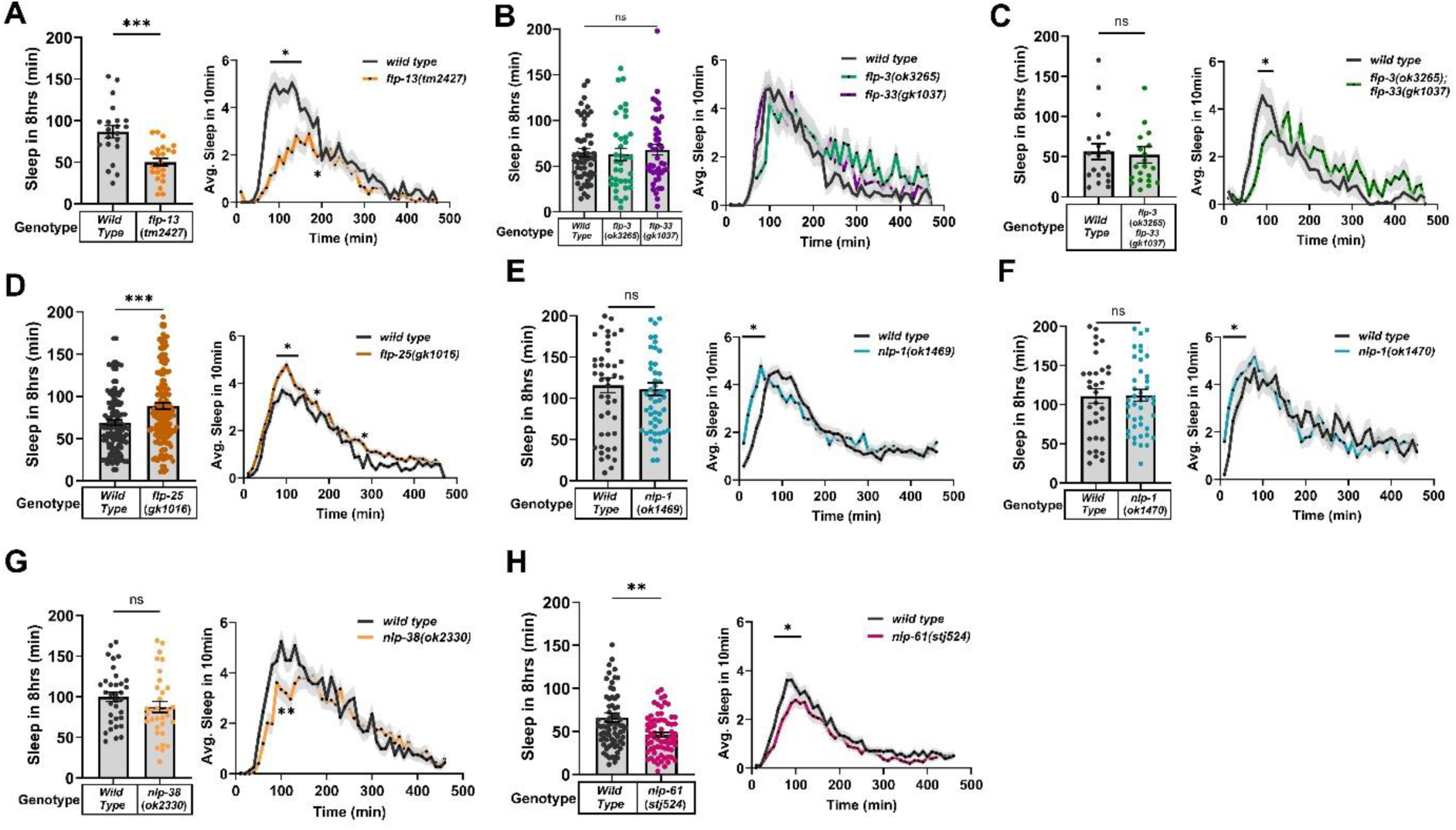
Multiple neuropeptide genes regulate stress-induced sleep. For each panel, the average total minutes of movement quiescence is displayed on the left, and the average minutes of movement quiescence in 10-minute windows on the right. **A)** Quiescence in wild-type (N=21) and *flp-13*(*tm2427*) (N=23) animals. Right panel – *p<0.05 at 70-140, and 190 minutes. **B)** Quiescence in wild-type (N=53), *flp-3*(*ok3265*) (N=37), and *flp-33*(*gk1037*) (N=44) animals. **C)** Quiescence in wild-type (N=19) and *flp-3*(*ok3265*); *flp-33*(*gk1037*) (N=21) animals. Right panel – *p<0.05 at 70-80 minutes. **D)** Quiescence in wild-type (N=124) and *flp-25*(*gk1016*) (N=171) animals. Right panel – *p<0.05 at 80-120, 170, and 280 minutes. **E)** Quiescence in wild-type (N=47) and *nlp-1*(*ok1469*) (N=53) animals. Right panel – *p<0.05 at 20-50 minutes. **F)** Quiescence in wild-type (N=34) and *nlp-1*(*ok1470*) (N=44) animals. Right panel – *p<0.05 at 20 minutes only. **G)** Quiescence in wild-type (N=34) and *nlp-38*(*ok2330*) (N=33) animals. Right panel – *p<0.05 at 80 and 100 minutes. **H)** Quiescence in wild-type (N=74) and *nlp-61*(*stj524*) (N=69) animals. Right panel – *p<0.05 at 60, 80, and 90 minutes. Student’s t-test was used for comparison between genotypes, and one-way ANOVA and Tukey’s test for three genotypes. Temporal comparisons were made using two-way ANOVA and Sidak’s test (*p<0.05, **p<0.01, ***p<0.001).

Ectopic OE of neuropeptide genes like *flp-11* and *flp-13* can induce sleep-like behavior in active, awake adults (Nelson *et al*. 2013; Nelson *et al*. 2014; Nath *et al*. 2016; Turek *et al*. 2016; Honer *et al*. 2020). To determine if *flp-3* and *flp-33* can also induce sleep, we used a heat shock promoter to drive their expression in all cells in wild-type animals. Two-hours following a mild heat stress (33°C), we manually measured body bends, and found that while *flp-13* strongly reduced locomotion, *flp-33* only mildly reduced it, and *flp-3* did not reduce it at all (**Figure S1A**). Thus, *flp-33* is capable of reducing activity, but neither gene can induce a sleep-like state. OE by transgenic multi-copy arrays can also affect sleep, like we have shown previously with *nlp-50*; transgenic lines carrying additional copies of the *nlp-50* gene displayed a strong reduction in SIS (Le *et al*. 2023). We made OE lines for *nlp-1* and *nlp-61*, and then measured UV-induced sleep, and found that *nlp-1* over expression did not alter SIS (**Figure S1B**), while *nlp-61* over-expression enhanced it (**Figure S1C**). Thus, *nlp-61* is required for sleep, and OE enhances it.

To determine if either *nlp-1* or *nlp-61* were required for developmentally-timed sleep (DTS), we measured L4 DTS in *nlp-1* mutants, but detected no differences from wild-type animals (**Figure S2A**), however, *nlp-61* mutants showed a significant reduction (**Figure S2B**). Thus, *nlp-61* is also required for DTS, and thus may play a broader role in sleep regulation.

### *nlp-1* and *nlp-61* are not endogenous ligands for the GPCR *npr-38*

The orphaned GPCR *npr-38* is required for SIS; loss of function of *npr-38* reduces sleep and disrupts avoidance behavior, while *npr-38* OE profoundly shifts the timing of sleep to an earlier period (Le *et al*. 2023). Where in the nervous system *npr-38* is functioning to coordinate the timing of sleep is unknown. *npr-38* is expressed in all of the cells of this circuit (**Figure 1**), except AIY (Taylor *et al*. 2021; Le *et al*. 2023). Cell-specific rescue of *npr-38* in some of the cells has been examined (RIS, RMG, ALA, AIB, ADL, and DVA), however, only expression in the ADL restored sleep, but did not alter the timing of sleep (Le *et al*. 2023). To test the other cells in this circuit, we restored *npr-38* in the *npr-38*(*stj367*) background in the RIF (Promoter: *nlp-6*), and RIM (Promoter: *cex-1*), and measured UV-induced sleep. RIF expression did not affect sleep in any noticeable manner (**Figure S3A**), while expression in RIM restored sleep amounts, but did not alter the timing of sleep (**Figure S3B**). These data indicate that *npr-38* can function in ADL and RIM to regulate sleep amounts, but how it coordinates the temporal aspects of sleep are still unknown.

OE phenotypes of neuropeptides require their downstream receptors, like what was observed with *flp-13* and *dmsr-1* (Iannacone *et al*. 2017), and phenotypes caused by multi-copy OE of GPCRs require the presence of endogenous ligands. This was demonstrated with the GPCR *egl-6* and ligands *flp-10*/*flp-17* in the egg-laying circuit (Ringstad and Horvitz 2008), and the GPCR *frpr-4* and *flp-13* during body bend control (Nelson *et al*. 2015). *nlp-1* and/or *nlp-61* were considered candidate ligands for *npr-38* (Personal Communication. Isabel Beets). So, we predicted that if *npr-38* would be required for the enhanced sleep we observed following *nlp-61* OE (**Figure S1C**), and that *nlp-61* would be required for the *npr-38* OE phenotypes. We found that *nlp-61* OE animals did indeed sleep significantly less in the absence of *npr-38*, compared to controls (**Figure S3C**). However, when *npr-38* OE in the *nlp-61*(*stj524*) background did not affect the temporal pattern of sleep (**Figure S3D**). These data do not support *npr-38* functioning as a receptor for *nlp-61*, but instead suggest that *npr-38* functions downstream of *nlp-61*. Next, we measured sleep in animals that OE *npr-38* in the *nlp-1*(*ok1469*) background and found a modulation of the sleep phenotype but not a requirement for the *nlp-1* peptides. Specifically, animals slept early, but the amount of sleep was higher in *nlp-1* mutants (**Figure S3E**). What these genetic interactions indicate at the mechanistic level is not clear, however, these data do not support *nlp-1* function as ligands for *npr-38*.

### *nlp-1* mutants display timing defects during acute heat exposure

*npr-38* OE animals display behavioral quiescence rapidly when exposed to acute heat stress, significantly earlier than wild-type controls (Le *et al*. 2023) (**Figure S3F**). We found that *nlp-1* mutants displayed early UV-induced sleep (**Figure 2E**), and in addition displayed behavioral quiescence faster on heat (**Figure S3G**). Taken together, we propose that *npr-38* and *nlp-1* coordinate the timing of sleep and our data indicate that they may function in parallel.

### Multiple GPCR genes are required for stress-induced sleep

Numerous neuropeptide GPCR genes (Janssen *et al*. 2010) are expressed in this circuit, which include FMRFamide peptide receptor (*frpr*), and neuropeptide receptor (*npr*) genes (**Figure 1**), of which we focused on highly expressed ones (Taylor *et al*. 2021). We first measured UV-sleep in *dmsr-1* (expressed in AIY and RIS) mutants, well-established sleep-defective animals (Iannacone *et al*. 2017). *dmsr-1*(*qn45*) mutants displayed severely reduced SIS compared to wild-type controls, yet the timing of sleep appeared to be mostly unaltered (**Figure 3A**). Next, we measured sleep in *frpr-4* (Expressed in RIS); our previous work demonstrated that *frpr-4* is a *flp-13* receptor, and that *frpr-4* OE induces behavioral quiescence (Nelson *et al*. 2014; Nelson *et al*. 2015). However, *frpr-4* deletion mutants (allele *ok2376*), did not display significant defects in heat-induced sleep (Nelson *et al*. 2015). Here, we measured UV-induced sleep, and surprisingly, *frpr-4*(*ok2376*) animals displayed a significant increase in sleep, with no changes in timing (**Figure 3B**). Considering our prior focus on *frpr-4*, we constructed a second loss-of-function mutant using CRISPR, *frpr-4*(*stj21*), and again found that sleep was significantly enhanced (**Figure 3C**). Thus, *frpr-4* signaling may function to negatively regulate sleep maintenance, but not timing.

**Figure 3:**
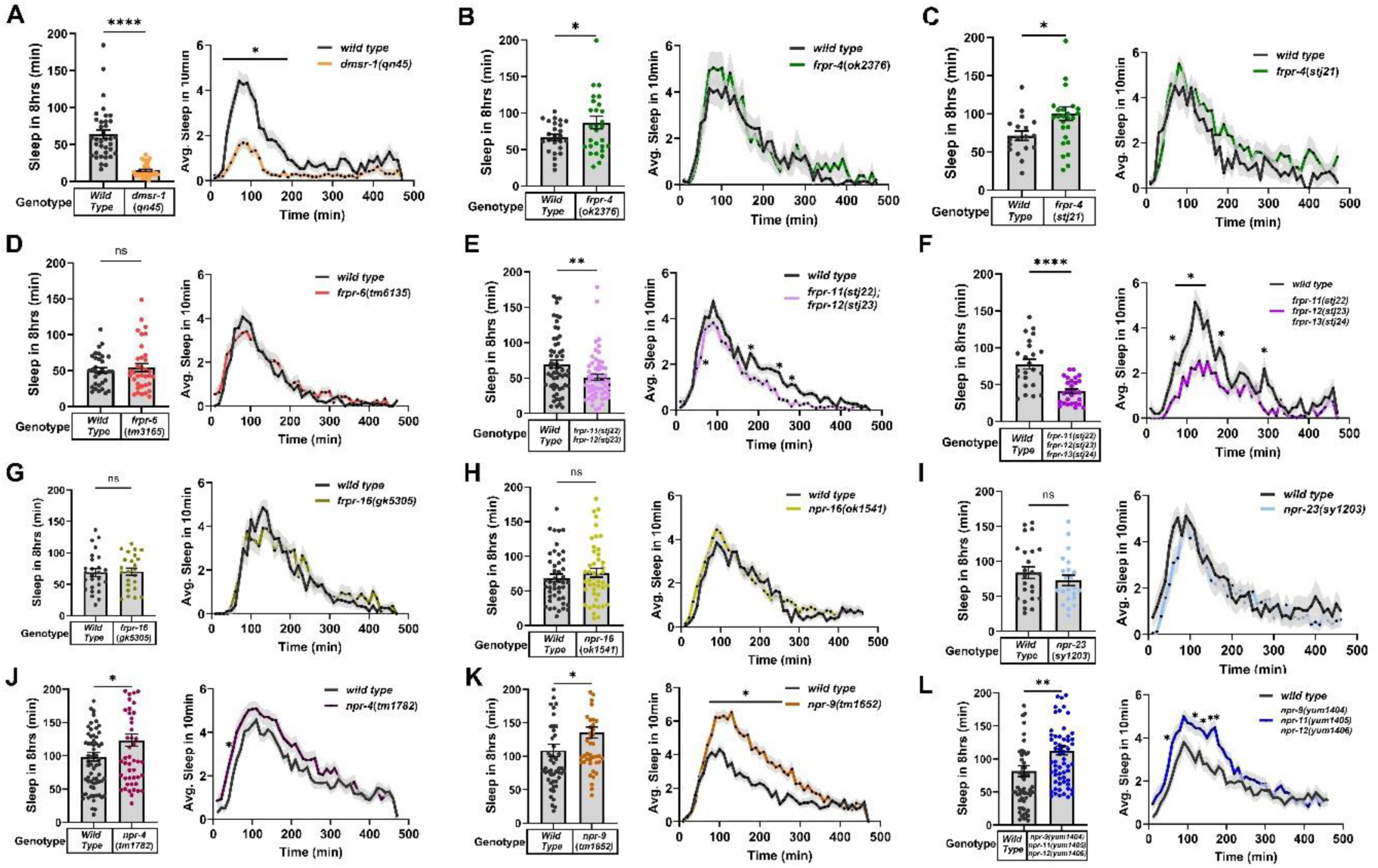
Multiple GPCR genes regulate stress-induced sleep. For each panel, the total minutes of movement quiescence (left), and average minutes of movement quiescence in 10-minute windows (right) is displayed. **A)** Quiescence in wild-type (N=36) and *dmsr-1*(*qn45*) (N=36) animals. Right panel – *p<0.05 at 40-170 minutes. **B)** Quiescence in wild-type (N=26) and *frpr-4*(*ok2376*) (N=29) animals, and **C)** wild-type (N=18) and *frpr-4*(*stj21*) (N=27) animals. **D)** Quiescence in wild-type (N=32) and *frpr-6*(*tm3165*) (N=35) animals. **E)** Quiescence in wild-type (N=59) and *frpr-11*(*sjt22*); *frpr-12*(*stj23*) (N=57) animals. Right panel – *p<0.05 at 60,180,250, and 280 minutes. **F)** Quiescence in wild-type (N=25) and *frpr-11*(*sjt22*); *frpr-12*(*stj23*); *frpr-13*(*stj24*) (N=28) animals. Right panel – *p<0.05 at 70,100-150,180, and 290 minutes. **G)** Quiescence in wild-type (N=26) and *frpr-16*(*gk5305*) (N=24) animals. **H)** Quiescence in wild-type (N=46) and *npr-16*(*ok1541*) (N=46) animals. **I)** Quiescence in wild-type (N=24) and *npr-23*(*sy1203*) (N=22) animals. **J)** Quiescence in wild-type (N=69) and *npr-4*(*tm1782*) (N=53) animals. Right panel – *p<0.05 at 40 minutes. **K)** Quiescence in wild-type (N=52) and *npr-9*(*tm1652*) (N=48) animals. Right panel – *p<0.05 at 90-260 minutes. **L)** Quiescence in wild-type (N=62) and *npr-9(yum1404); npr-11(yum1405)*; *npr-12(yum1406)* (N=65) animals. Right panel – *p<0.05 at 60,120,14,160-170 minutes. Student’s t-test was used to statistically compare the total quiescence of two genotypes (*p<0.05, **p<0.01, ****p<0.0001), and temporal comparisons between genotypes were made using two-way ANOVA followed by Sidak’s test.

*frpr-6* (Expressed in AIY, RIS, and RMG), *frpr-11* (Expressed in AIY and RIF), *frpr-12* (Expressed in AIB and AIY), *frpr-13* (Expressed in AIY, DVA, and RIF) and *frpr-16* (Expressed in RIM and RIS) are potential paralogs with *frpr-4*, and expressed in this circuit. *frpr-6* is the most similar to *frpr-4*, however, deletion mutants did not display sleep defects (**Figure 3D**). *frpr-11*, *frpr-12*, and *frpr-13* are highly similar to one another, and located adjacently on chromosome V, with *frpr-11* and *frpr-12* being almost identical, indicating they likely arose by gene duplication (Mccoy *et al*. 2014). We constructed an *frpr-11*; *frpr-12* double mutant, and measured a modest but significant reduction in total levels of SIS, as well as at various time points throughout sleep (**Figure 3E**). Next, we generated an *frpr-11*; *frpr-12*; *frpr-13* triple mutant, and found that sleep was even more reduced (**Figure 3F**), with the timing mostly unaltered. These data indicate that *frpr-11*, *frpr-12*, and *frpr-13* function redundantly to positively regulate sleep maintenance.

Next, we measured sleep in *frpr-16*, *npr-16* (Expressed in ADL, AIB, AIY, RIF, and RIS), and *npr-23* (Expressed in AIB, AIY, ALA, DVA, RIF, RIS, and RMG) mutants, however, did not detect any changes in sleep (**Figure 3G-I**). However, when we measured sleep in *npr-4* (Expressed in ADL, AIB, RIF, RIM, RIS, and RMG), and *npr-9* (Expressed in AIB, ALA, and RIM) mutants, sleep was significantly enhanced. Total sleep was increased in both mutants, sleep occurred slightly earlier in *npr-4* animals, while sleep was enhanced at numerous time-points during the normal sleep cycle in *npr-9* mutants (**Figure 3J, K**). We also measured sleep in an available *npr-9*;*npr-11*;*npr-12* triple mutant (*npr-11* is expressed in ADL, AIB, AIY, and RMG; *npr-12* is expressed in DVA, RIF, and RIS) (Pu *et al*. 2023), and again, sleep was enhanced, and in this case shifting slightly earlier (**Figure 3L**). Total levels did not appear to be higher than *npr-9* single mutants. Overall, these data suggest that *npr-4* and *npr-9* negatively regulate sleep maintenance, and potentially sleep timing. More broadly, numerous GPCRs may modulate sleep in unique and opposing ways, indicating complex antagonistic signaling mechanisms modulate sleep maintenance and timing.

### The two-pored potassium channel *twk-16* negatively regulates sleep

The gene *twk-16*, which codes a mechanically gated, two-pored weakly-inward rectifying (TWIK) potassium channel (Salkoff *et al*. 2001; Das *et al*. 2021), is selectively expressed in the DVA (Puckett robinson *et al*. 2013). *twk-16* may regulate DVA membrane potential which regulates body wall muscles, and bending angle (Das *et al*. 2021). The role for DVA during sleep regulation is unclear, however, activation of a red-light induced adenylyl cyclase called IlaC22 (Ryu *et al*. 2014), in the DVA impairs SIS (Cianciulli *et al*. 2019), suggesting it is wake promoting. To further explore this, we generated two loss-of-function alleles, *stj13* and *stj16* (**Figure 4A**). Both *twk-16*(*stj13*) and *twk-16*(*stj16*) animals displayed modest but significant enhancement of sleep, with the peak of sleep between 90-120 minutes post-stress being consistently higher than controls (**Figure 4B, C**), however the timing of sleep was unaffected. Next, we made a transgenic multi-copy rescue strain, in which wild-type *twk-16* genes driven from their endogenous promoter, were expressed in the *twk-16*(*stj13*) background. This OE rescue of *twk-16* caused a significant impairment of total sleep (**Figure 4D**). These data suggest that *twk-16* negatively regulates sleep maintenance, possibly by maintaining resting potential of the wake-promoting DVA neuron.

**Figure 4:**
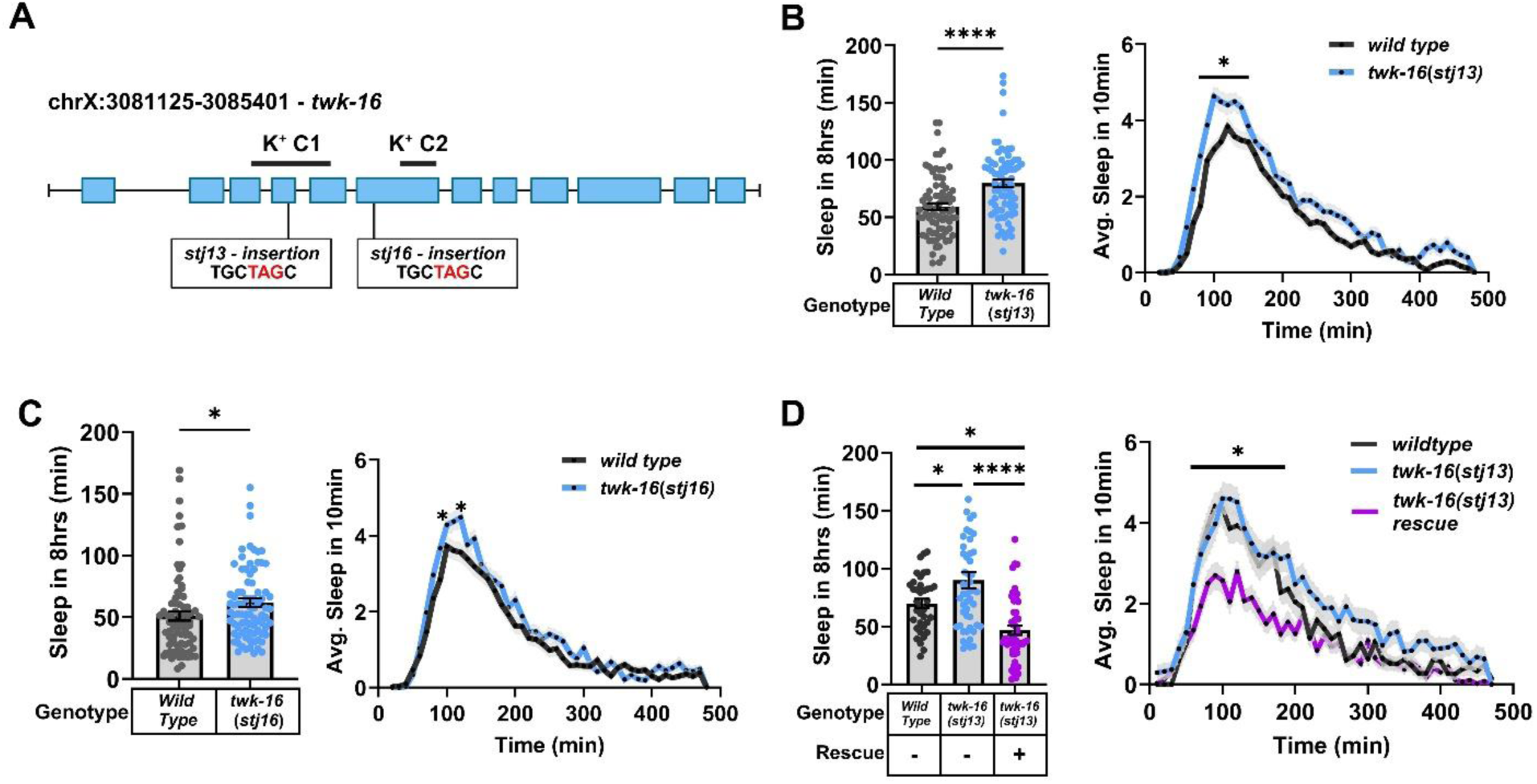
*twk-16* loss-of-function enhances sleep. **A)** Schematic of the gene and allele structure for the two-pored potassium channel gene *twk-16*. K^+^C1 = Potassium Channel 1, and K^+^C2 = Potassium Channel 2. **B)** Quiescence in wild-type (N=77) and *twk-16*(*stj13*) (N=76) animals. Right panel – *p<0.05 at 60-100, and 120 minutes. **C)** Quiescence in wild-type (N=79) and *twk-16*(*stj16*) (N=84) animals. Right panel – *p<0.05 at 80 and 120 minutes. **D)** Quiescence in wild-type (N=37), *twk-16*(*stj13*) (N=51), and *twk-16*(*stj13*) transgenic animals expressing wild-type *twk-16* copies (N=51). Right panel – Comparing *twk-16*(*stj13*) to rescued strain, *p<0.05 at 60-230 minutes. Student’s t-test was used to statistically compare two genotypes, one-way ANOVA followed by Tukey’s test to compare three genotypes, and temporal comparisons were made using two-way ANOVA followed by Sidak’s test (*p<0.05, ****p<0.0001).

### *eat-4* and *glr-5* are required for stress-induced sleep maintenance and timing

The vesicular glutamate transporter gene *eat-4* (Lee *et al*. 1999) (Expressed in ADL, AIB, DVA, and RIM) and the non-NMDA glutamate receptor *glr-5* (Brockie *et al*. 2001) (Expressed in AIB, ALA, DVA, RIF, RIM, RIS, and RMG) are expressed in a number of cells in this circuit (**Figure 1**). So, we measured sleep in *eat-4*(*ky6*) loss-of-function mutants (Ohnishi *et al*. 2011), and found that SIS was reduced (**Figure 5B**), suggesting that glutamate positively regulates SIS. Next, we used CRISPR to tag the 3’ end of *eat-4* with a degron sequence (*stj606* allele), to allow for the degradation of EAT-4 protein using the auxin-inducible degradation (AID) system (Zhang *et al*. 2015). To determine the usefulness of this strain, we first measured feeding rate (i.e., pharyngeal pumping), a phenotype known to be impaired in *eat-4* loss-of-function mutants (Lee *et al*. 1999; Dalliere *et al*. 2016). We first measured pumping in wild-type, *eat-4*(*ky5*), *eat-4*(*ad450*) mutants, and *eat-4*(*stj606*) animals in the absence of auxin. Pumping was significantly reduced in all 3 mutants, however, *eat-4*(*stj606*) animals pumped significantly more than the other known loss-of-function strains (**Figure S4A**). We concluded that the 3’degron was likely interfering with the function of *eat-4*. To determine if *eat-4* protein degradation was possible, we expressed the plant gene *tir1* (Zhang *et al*. 2015), in all cells using the *eft-3* promoter. Auxin treatment for 5 or 24 hours did not further reduce pumping (**Figure S4B**). We conclude that *stj606* is a hypomorphic allele and not useful for the AID system. So, we measured sleep in *eat-4*(*stj606*) animals in the absence of auxin, and found sleep to be reduced but appeared less severe than *eat-4*(*ky5*) animals (**Figure 5C**). Taken together, these data suggest that glutamate signaling positively regulates SIS.

**Figure 5:**
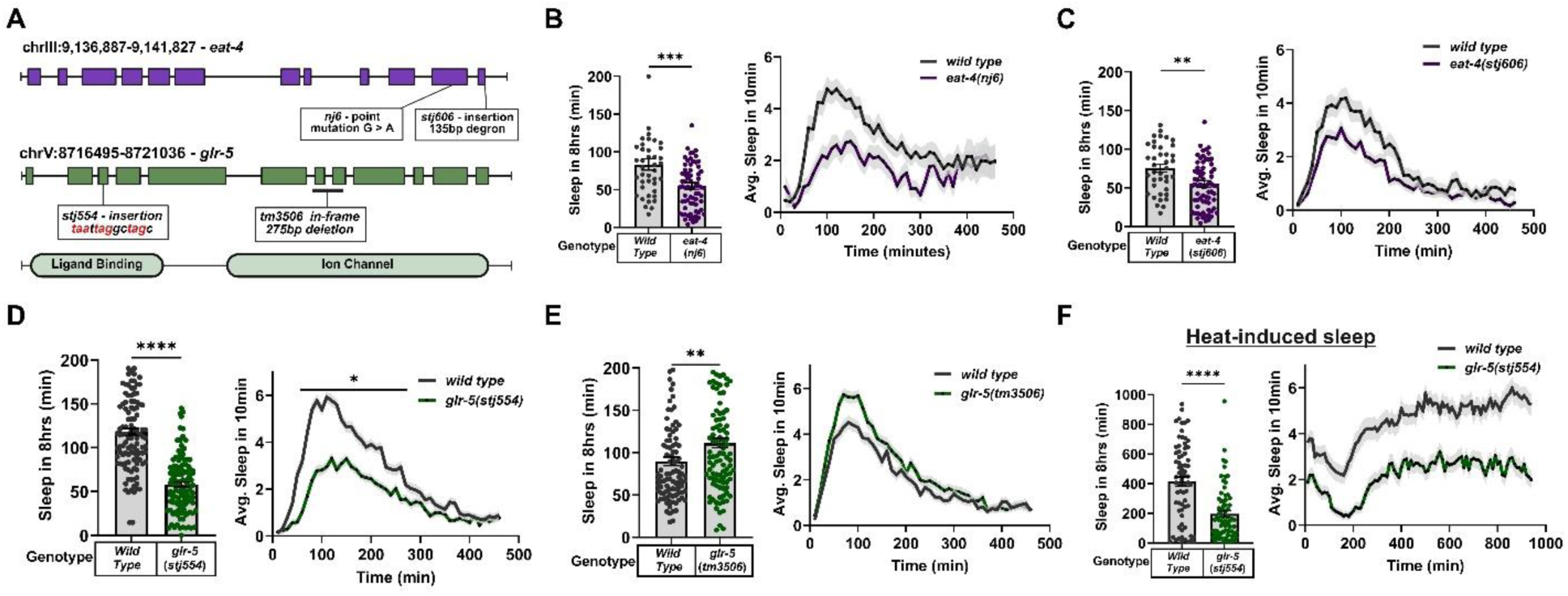
Glutamate signaling is required for stress-induced sleep. **A)** Schematic of the gene and allele structure for *eat-4* and *glr-5*. For **B-F**, the total minutes of movement quiescence (left), and average minutes of movement quiescence in 10-minute windows (right) is displayed. **B)** Quiescence during UV-induced SIS in wild-type (N=46) and *eat-4*(*nj6*) (N=31) animals. **C)** Quiescence during UV-induced SIS in wild-type (N=40) and *eat-4*(*stj606*) (N=71) animals. **D)** Quiescence during UV-induced SIS in wild-type (N=113) and *glr-5*(*stj554*) (N=121) animals. Right panel – *p<0.05 40-240 minutes, two-way ANOVA followed by Sidak’s test. **E)** Quiescence during UV-induced SIS in wild-type (N=99) and *glr-5*(*tm3506*) (N=105) animals. **F)** Quiescence during heat-induced SIS in wild-type (N=113) and *glr-5*(*stj554*) (N=121) animals. Student’s t-test was used to statistically compare sleep totals (**p<0.01, ***p<0.001, ****p<0.0001).

Next, we measured sleep in two *glr-5* mutants, *glr-5*(*tm3506*), which contain a small in-frame deletion, and *glr-5*(*stj554*), who carry an insertion of 3 stop codons in each reading frame in the third exon of the gene, presumably a null (**Figure 5A**). We found that *glr-5*(*stj554*) animals displayed impaired sleep; it was initiated later, and lower overall (**Figure 5D)**. These data suggest that *glr-5* promotes the initiation and maintenance of sleep. Considering that UV-sleep was so impaired in these animals, we wanted to test if *glr-5* was required for other forms of sleep. Heat-induced sleep, and L4 developmentally timed-sleep were also reduced in these mutants (**Figure 5F and Figure S2D**). Additionally, we measured the acute response to noxious heat, and found that animals displayed increased activity compared to controls (**Figure S5A, B)**. We also assessed their response to directed blue light, a noxious stimulant known to initiate avoidance (Edwards *et al*. 2008), and found that *glr-5* mutants responded slower than controls **(Figure S5C)**. Thus, *glr-5* is required for behaviors of the stress response, including sensory responsiveness to noxious stimuli, sleep timing, and sleep maintenance.

*glr-5*(*tm3506*) mutants have egg laying defects (Wen *et al*. 2020), and an impaired response to chemical repellants (Zou *et al*. 2018). Surprisingly, UV-induced sleep was modestly but significantly enhanced in these animals (**Figure 5E**). The *tm3506* allele is a 275bp deletion which likely truncates an ion channel domain by 43 amino acids (**Figure 5A**), so may not be loss of function. To confirm this enhanced sleep phenotype, we analyzed *glr-5*(*stj554*) and *glr-5*(*tm3506*) animals, side-by-side in the same experiments. As expected, *glr-5*(*stj554*) animals displayed reduced sleep, while *glr-5*(*tm3506*) animals slept significantly more (**Figure S6A**). We conclude that *tm3506* disrupts *glr-5* function, however, in the context of sleep regulation this results in a gain-of-function phenotype. What this means mechanistically is still unclear.

### *glr-5* functions in the ALA, RIS, AIB, and RIM to control sleep timing and amounts

We next sought to determine in which cells *glr-5* functions during sleep. We constructed transgenic *glr-5*(*stj554*) animals that expressed the wild-type *glr-5* from various promoters expressed in AIB (*npr-9*), ALA (*ver-3*), DVA (*twk-16*), RIM (*cex-1*), RIS (*flp-11* or *aptf-1*), and RMG (*flp-21*) (Popovici *et al*. 2002; Kim and Li 2004; Van Buskirk and Sternberg 2010; Puckett robinson *et al*. 2013; Turek *et al*. 2013; Campbell *et al*. 2016; Taylor *et al*. 2021), as well as from the *glr-5* endogenous promoter (genomic rescue). Genomic rescue of *glr-5* increased sleep close to wild-type levels (**Figure 6A**). We speculate that the lack of full rescue could be due to dosage effects of the multi-copy array. Rescuing *glr-5* in the ALA and RIS, RMG, AIB, or DVA did not restore sleep (**Figure 6B-E**). In fact, expression in the ALA and RIS further reduced sleep lower than *glr-5* mutants alone (**Figure 6B**). However, there was a striking rescue of the temporal defects of *glr-5* mutants (**Figure 6H**). These data suggest that sleep initiation (i.e., timing) is regulated by *glr-5* through the RIS and/or ALA. The RIS is highly connected to the AIB and RIM interneurons by gap and chemical junctions (**Figure 1**) (White *et al*. 1986). Rescue in the RIM modestly increased sleep levels but did not alter the timing of sleep (**Figure 6F**), however, expression in the AIB, RIM, and RIS restored sleep to wild-type levels, and rescued the timing defects (**Figure 6G and 6I**). We propose a model in which *glr-5* functions to regulate the sleep timing through the RIS, and maintenance through AIB, RIS, and RIM, which form a highly connected 3-celled circuit (**Figure S7**).

**Figure 6:**
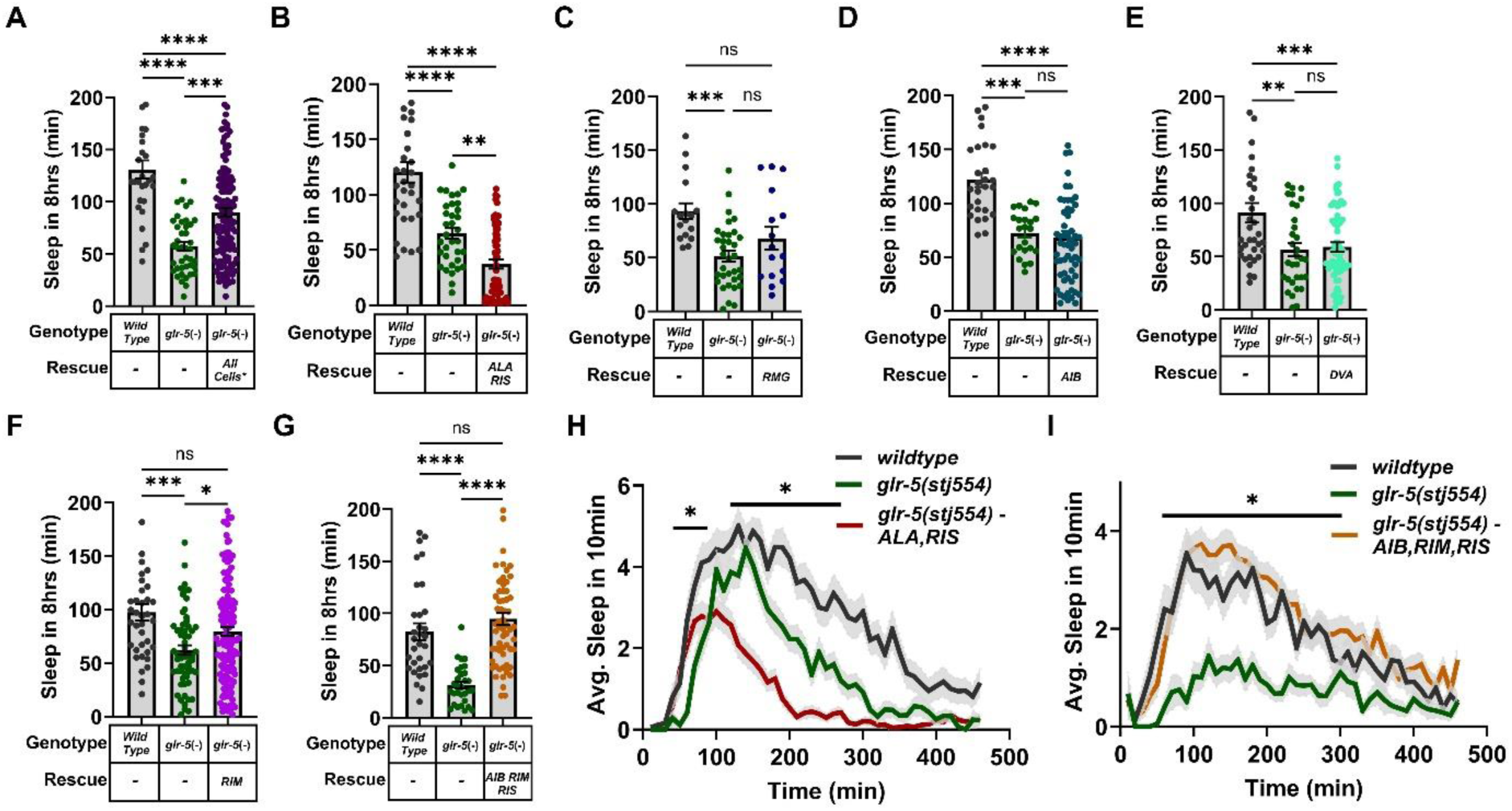
*glr-5* functions in the AIB, RIM, and RIS to regulate UV-induced sleep. **A)** Quiescence during UV-induced SIS in wild-type (N=29), *glr-5*(*stj554*) (N=41), and transgenic *glr-5*(*stj554*) (N=68) (Rescue: *All cells in which *glr-5* is expressed; Promoter: *glr-5*) animals. **B)** Quiescence during UV-induced SIS in wild-type (N=32), *glr-5*(*stj554*) (N=34), and transgenic *glr-5*(*stj554*) (N=65) (Rescue: ALA and RIS; Promoters: *ver-3* and *flp-11*) animals. **C)** Quiescence during UV-induced SIS in wild-type (N=17), *glr-5*(*stj554*) (N=34), and transgenic *glr-5*(*stj554*) (N=16) (Rescue: RMG; Promoter: *flp-21*) animals. **D)** Quiescence during UV-induced SIS in wild-type (N=27), *glr-5*(*stj554*) (N=23), and transgenic *glr-5*(*stj554*) (N=58) (Rescue: AIB; Promoter: *npr-9*) animals. **E)** Quiescence during UV-induced SIS in wild-type (N=35), *glr-5*(*stj554*) (N=32), and transgenic *glr-5*(*stj554*) (N=63) (Rescue: DVA; Promoter: *twk-16*) animals. **F)** Quiescence during UV-induced SIS in wild-type (N=36), *glr-5*(*stj554*) (N=30), and transgenic *glr-5*(*stj554*) (N=60) (Rescue: RIM; Promoter: *cex-1*) animals. **G)** Quiescence during UV-induced SIS in wild-type (N=32), *glr-5*(*stj554*) (N=30), and transgenic *glr-5*(*stj554*) (N=61) (Rescue: AIB, RIM, RIS; Promoters: *aptf-1* and *cex-1*) animals. For **A-G**, statistical significance was calculated by one-way ANOVA followed by Tukey’s test (**p<0.01, ***p<0.001, ****p<0.0001). **H)** Average quiescence in 10-minute windows in wild-type (N=32), *glr-5*(*stj554*) (N=34), and transgenic *glr-5*(*stj554*) (N=65) (Rescue: ALA and RIS; Promoters: *ver-3* and *flp-11*) animals. **I)** Average quiescence in 10-minute windows in wild-type (N=32), *glr-5*(*stj554*) (N=30), and transgenic *glr-5*(*stj554*) (N=61) (Rescue: AIB, RIM, RIS; Promoters: *aptf-1* and *cex-1*) animals. Comparisons were made using two-way ANOVA followed by Sidak’s test (*p<0.05).

## Discussion

Transitions between sleep and arousal are coordinated by antagonistic action of sleep- and wake-active interneurons, which orchestrate global brain states (Nichols *et al*. 2017). Sleep is dominated by the action of GABAergic and neuropeptidergic sleep neurons (Saper *et al*. 2010; Bringmann 2018). Their activation during daily sleep of insects and mammals, and developmentally-timed sleep of nematodes, is influenced by circadian input, and homeostasis (Saper *et al*. 2005; Monsalve *et al*. 2011). This regulation is coordinated by transcriptional-translational feedback loops of conserved circadian clock genes. As homeostatic sleep drive builds throughout the day, one function of the clock may be to ensure that arousal is maintained and sleep initiation is repressed until an animal can safely rest (Shafer and Keene 2021). If sleep pressures are enhanced outside of normal daily conditions, such as by cytokine release from immune cells, or diffusible factors from damaged cells, this can override circadian control and induce sleep during the day, or heighten it at night (Imeri and Opp 2009; Davis and Raizen 2017). Stress-induced sleep of invertebrates has become an important model for exploring the mechanisms of sleep outside of circadian regulation, but also for understanding how sleep and arousal are balanced by organisms (Hill *et al*. 2014; Lenz *et al*. 2015).

Mechanisms have likely evolved to negatively and positively regulate sleep initiation, and sleep depth (i.e., maintenance), considering selective evolutionary pressures exist for and against sleep. Sleep is essential, yet renders animals vulnerable to predation, and limits foraging and reproduction (Keene and Duboue 2018; Anafi *et al*. 2019). The opposing mechanisms that underlie sleep and arousal, both at the genetic and cellular level need further study. Here, we used *C. elegans* stress-induced sleep as a model to identify new signaling molecules that regulate sleep timing and maintenance.

Upon threat exposure, *C. elegans* will initiate stereotypical avoidance responses, consisting of accelerated movement and heightened arousal (Culotti and Russell 1978). Injury requires recovery sleep for survival; however, sleep must be precisely timed in order to prevent further damage. The ALA and RIS neurons are required for sleep, their depolarization induces sleep, and their activation correlates with sleep in real-time (Turek *et al*. 2013; Hill *et al*. 2014; Nelson *et al*. 2014; Konietzka *et al*. 2020). As with sleep in other animals, opposing mechanisms must be in place to coordinate avoidance, sleep, and arousal. Our work identified the molecules *nlp-14*(expressed in ALA) and *npr-38* as candidates for coordinating these behaviors. *nlp-14* mutants display early sleep (Honer *et al*. 2020), as do *npr-38* OE animals (Le *et al*. 2023). The *nlp-14* receptor, and *npr-38* ligands, as well as an understanding of the timing mechanisms are unknown. This motivated our current study, where we sought to identify new sleep-regulatory genes expressed in ALA, RIS, and/or a subset of cells connected with them (**Figure 1**). Using this approach, we identified signaling molecules that both positively and negatively regulate sleep timing, and sleep maintenance, which will allow for future mechanistic studies using the stress-induced sleep model of *C. elegans*.

### Sleep timing

The timing of sleep neuron activation is likely a crucial component of the temporal dynamics of these events. During the avoidance phase, molecules that suppress sleep initiation will dominate, however, as cells incur damage they will release *siss-1* epidermal growth factors which signal to the ALA and RIS, through the *let-23* receptor and second messenger pathways controlled by phospholipase C (Van Buskirk and Sternberg 2007; Hill *et al*. 2024). Controlled doses of UV irradiation, can induce sleep that is highly experimentally reproducible, such that animals remain active for ∼30-45 minutes prior to sleep induction (Debardeleben *et al*. 2017). Genetic manipulations that cause sleep to occur earlier involve negative regulators of sleep induction, while those that shift sleep to later time points are positive regulators. In this current study we have identified both. *nlp-1* mutants initiate sleep early, thus *nlp-1* peptides may act as negative regulators of sleep induction. They are expressed in the RIM which is presynaptic to RIS, thus could function to inhibit RIS as sleep drive increases following injury. We propose a model in which *nlp-1* functions in parallel with *nlp-14* and *npr-38* to prevent sleep initiation during the avoidance phase (**Figure S7A**). Loss of *flp-3* and *flp-33*, *flp-13*, or *glr-5* initiates sleep later, thus they represent positive regulators. Even in the context of these mutations, sleep eventually initiated, presumably via *siss-1*-mediated activation of ALA and RIS. However, we again propose a model where multiple signals function at the level of sleep neurons to allow for robust activation at the correct time (**Figure S7A**). Our rescue data, in particular, indicate that *glr-5* functions directly on RIS to positively regulate sleep initiation (**Figure 6H**). Future work will be focused on identify the sites of action for the other molecules identified here.

### Sleep maintenance

Multiple mutations that enhance or impair sleep in *Drosophila* (Shafer and Keene 2021), mammals (Shen *et al*. 2022), and *C. elegans* (Honer and Nelson 2024) have been identified, indicating complex genetic redundancy regulates depth (i.e., maintenance) of sleep. During stress-induced sleep of *C. elegans*, the RIS sleep neuron itself can in part regulate the duration and depth of sleep, by inhibitory autocrine signaling by *flp-11*/*dmsr-1* (Rossi *et al*. 2025). While other peptides released from ALA and RIS are believed to contribute downstream of the sleep neurons. Here, we measured sleep in signaling molecules expressed in cells that upstream of ALA and RIS, thus may play more of a role in the maintenance of sleep neuron activation. The molecules identified here who may function as positive regulators of sleep maintenance include *nlp-38*, *nlp-61*, *flp-3*, *flp-33*, *frpr-11*, *frpr-12*, *frpr-13*, *eat-4*, and *glr-5*. The potential negative regulators are *twk-16*, *frpr-4*, *flp-25*, *npr-4*, and *npr-9* (**Figure S7A**). The list of genes that regulate sleep, even in simple animals like *C. elegans*, continues to grow, and suggests that throughout evolution, multiple and potentially redundant opposing mechanisms have arisen. As with timing, it will be important to determine on which cells these molecules function and the underlying mechanisms.

### Glutamate signaling and sleep

Glutamatergic transmission is conserved across phylogeny, and both positively and negatively regulates circadian sleep in *Drosophila* and mammals (John *et al*. 2008; Zimmerman *et al*. 2008; Guo *et al*. 2016; Yoon *et al*. 2025). In *C. elegans*, glutamate signaling via the *glr-2* receptor promotes arousal during developmentally-timed sleep (Choi *et al*. 2015), and here, we identified that glutamate signaling via *glr-5* positively regulates stress-induced sleep. Interestingly, *glr-5* acts directly on the RIS sleep neuron, as well as AIB and RIM which are gapped to RIS, to regulate sleep maintenance and timing. Numerous glutamatergic neurons are presynaptic to these cells, an important next step will be to identify how glutamate transmission communicates with sleep neurons (**Figure S7B**).

Some recent evidence support a conserved function for glutamate signaling, considering the *Drosophila* ortholog Eye-enriched kainate receptor (Ekar), was identified in a genetic screen as a sleep regulating gene (Stirtz *et al*. 2026). Also, the mammalian ortholog GRIK4 has been implicated with sleep regulation in mice. GRIK4-positive thalamic neurons are activated in response to arousal from slow-wave sleep (Shin *et al*. 2023). Thus, glutamate signaling through *glr-5*-like receptors, may be a common mechanism of sleep coordination, maintenance, and arousal across phylogeny.

## Methods

### Worm maintenance and strains

Animals were maintained at 20°Celsius on agar plates containing nematode growth medium and fed the OP50 derivative bacterial strain DA837 (Davis *et al*. 1995). The strains that were used in this study were generated in the Nelson lab, or obtained from the National BioResource Project (NBRP), the *Caenorhabditis* Genetics Center (CGC), or the Raizen Lab (University of Pennsylvania) (**Table S1)**.

### Molecular biology and transgenesis

To generate DNA for transgenesis, the polymerase chain reaction (PCR), or overlap extension-PCR (OE-PCR) was used (Nelson and Fitch 2011). Transgenic animals were made by microinjection (Stinchcomb *et al*. 1985). To make genomic rescue lines for *twk-16* and *glr-5*, and multi-copy over-expression lines for *nlp-1* and *nlp-61*, the 5’UTR, coding sequence, and 3’UTR were amplified by from wild-type genomic DNA for each corresponding gene. The *twk-16* wild-type amplicons were injected into *twk-16*(*stj13*) animals, and the *glr-5* wild-type PCR products were injected into *glr-5*(*stj554*) animals. PCR products of either *nlp-1* or *nlp-61* were injected into wild-type animals (N2). To generate cell-specific rescue constructs for *glr-5* and *npr-38*, and inducible over-expression constructs for *flp-3* and *flp-33*, OE-PCR was used. Constructs for cell-specific rescue were made by first amplifying the genomic promoter sequence of the genes *ver-3* (ALA) (Popovici *et al*. 2002; Van buskirk and Sternberg 2010), *flp-11* (RIS) (Turek *et al*. 2013), *npr-9* (AIB) (Campbell *et al*. 2016), *aptf-1* (AIB + RIS) (Turek *et al*. 2013; Taylor *et al*. 2021), *flp-21* (RMG) (Kim and Li 2004), *nlp-6* (RIF), and *cex-1* (RIM) (Taylor *et al*. 2021). The corresponding promoter was fused by OE-PCR to the coding sequence and 3’UTR of *glr-5* or *npr-38*. To generate *flp-3* and *flp-33* over-expression constructs, the promoter from the gene *hsp-16.2* was fused to the coding sequence and 3’UTR of the corresponding gene by OE-PCR. Oligonucleotides are listed in **Table S2**.

### Construction of mutants

The *nlp-61*, *frpr-4*, *frpr-11*, *frpr-12*, *frpr-13*, *twk-16* and *glr-5* loss-of-function alleles were generated using a defined CRISPR approach (Paix *et al*. 2017). Single-stranded oligonucleotides composed of an insertion sequence containing stop codons and a NheI (New England Biolabs) restriction site, flanked by 35bp homology arms were constructed (Integrated DNA Technologies (IDT)) (**Table S2**). For each CRISPR mutant, we simultaneously edited the *dpy-10* gene to allow for screening using the roller or dumpy physical phenotypes (Paix *et al*. 2017). A mixture of guide RNA (gRNA) duplexed with Alt-R® CRISPR-Cas9 tracrRNA (IDT), Alt-R® S.p. Cas9 Nuclease V3 (IDT), and oligonucleotide repair templates were injected into day-1 adult wild-type (N2) animals. For the *frpr-11*, *frpr-12*, and *frpr-13* mutants, we first made the *frpr-11*; *frpr-12* double mutant by injecting the CRISPR reagents for both genes into wild-type animals. Once these mutants were isolated, we then injected the *frpr-13* CRISPR mix into the double mutant. To identify all mutants, F1 animals displaying physical phenotypes (dumpy or roller) were singled, allowed to self, and screened by PCR followed by restriction digest. If the animals were homozygous for the *dpy-10* mutation they were crossed with N2. For the AID-tagged *eat-4* mutant, we first PCR amplified the AID sequence from the plasmid pLZ29 (Addgene) (Zhang *et al*. 2015), using two different primer pairs, one with 100bp homology arms and the other with 35bp arms, which are complementary to the same sequence at the 3’end of *eat-4*. These two PCR products were melted together to increase efficiency of insertion, as previously described (Ghanta and Mello 2020). The melted PCR product was used as the repair template to generate the strain SJU606. Candidate mutants were screened by PCR across the insertion site, and identified based on size. All alleles were confirmed by sanger sequencing (Genewiz).

### WorMotel behavioral assays

Movement quiescence for both DTS and SIS was measured using the WorMotel, as previously described (Churgin *et al*. 2017). For SIS, L4 animals were picked the day prior to the experiment, and allowed to grow to adulthood overnight. Day-1 adults were transferred to the agar surfaces of 24-welled Polydimethylsiloxane (PDMS) microchips, and exposed to either UV irradiation, or noxious heat, and then imaged for 8 hours. UV stress was applied by placing the entire microchip into a cross linker (Ultraviolet, 254 UVP), which was set to 1500 J/m^2^ of UV light, as described (Debardeleben *et al*. 2017). For heat stress, the microchip was placed in a petri dish which was sealed with parafilm, and submerged in a 37°C water bath for 30 minutes, and then imaged for 12 hours. For each experiment, wild-type, and the relevant mutant, and/or transgenic animals were run on the same PDMS chip over multiple trials. For DTS, wild-type and mutant animals were placed on microchips during the L4 stage prior to entering lethargus, and allowed to develop to young adulthood while being imaged over a 16-hour time period. For both DTS and SIS, quiescence was quantified using pixel subtraction, as previously described (Yu *et al*. 2014; Churgin *et al*. 2017), and total quiescence over the duration of the sleep cycles were compared between genotypes using Student’s t-test (Wild type and vs. a single mutant) or One-way ANOVA followed by Tukey’s multiple comparisons test (Wild type, and two other genotypes). Temporal patterns were compared by Two-way ANOVA followed by Sidak’s multiple comparisons test.

### Feeding rate of *eat-4* mutants

To measure pumping rate, day-1 adults were manually observed for 20-second intervals at 80X magnification using a stereomicroscope. A single pump was defined as one full movement of the pharyngeal grinder during both the contraction and relaxation phase (Raizen *et al*. 2012). For experiments involving auxin, the derivative indole-3-acetic acid (ThermoFisher) was supplemented in the bacterial food, as described (Zhang *et al*. 2015), and pumping was measured after 5 or 24 hours. Averages were calculated and compared by One-way ANOVA followed by Tukey’s multiple comparisons test.

### Acute quiescence upon heat exposure

We used manual observations to quantify the acute quiescence response to heat. For each experiment, 10 day-1 adults were transferred to standard growth plates, which were placed on a slide warmer set at 37°C for 30 min. The number of animals that were quiescent for both moving and feeding was counted every 1, 5, or 10 minutes, depending on the experiment. This was repeated over multiple trials. The fraction of quiescence was calculated by dividing the number of quiescent worms by the total number of worms, and then averaged across trials. Comparisons were made using two-way ANOVA followed by Sidak’s multiple comparisons.

### Avoidance response to blue light

To quantify blue light avoidance, body bends were counted manually over a 1-min span in day-1 adult animals, in the presence of directed blue light using a fluorescent stereomicroscope (Leica). A body bend was defined as one movement in either direction, in which the head changed direction. The average number of bends were calculated and compared across genotypes using one-way ANOVA followed by Tukey’s multiple comparisons test.

## Acknowledgements

Some strains were provided by the CGC, which is funded by NIH Office of Research Infrastructure Programs (P40 OD010440). We would also like to thank the SJU Summer Scholars Program and the John P. McNulty Fellows Program. This work was supported by the National Science Foundation (IOS 1845020 and IOS 2445004 to MDN).

## Supporting Information

**Figure S1:**
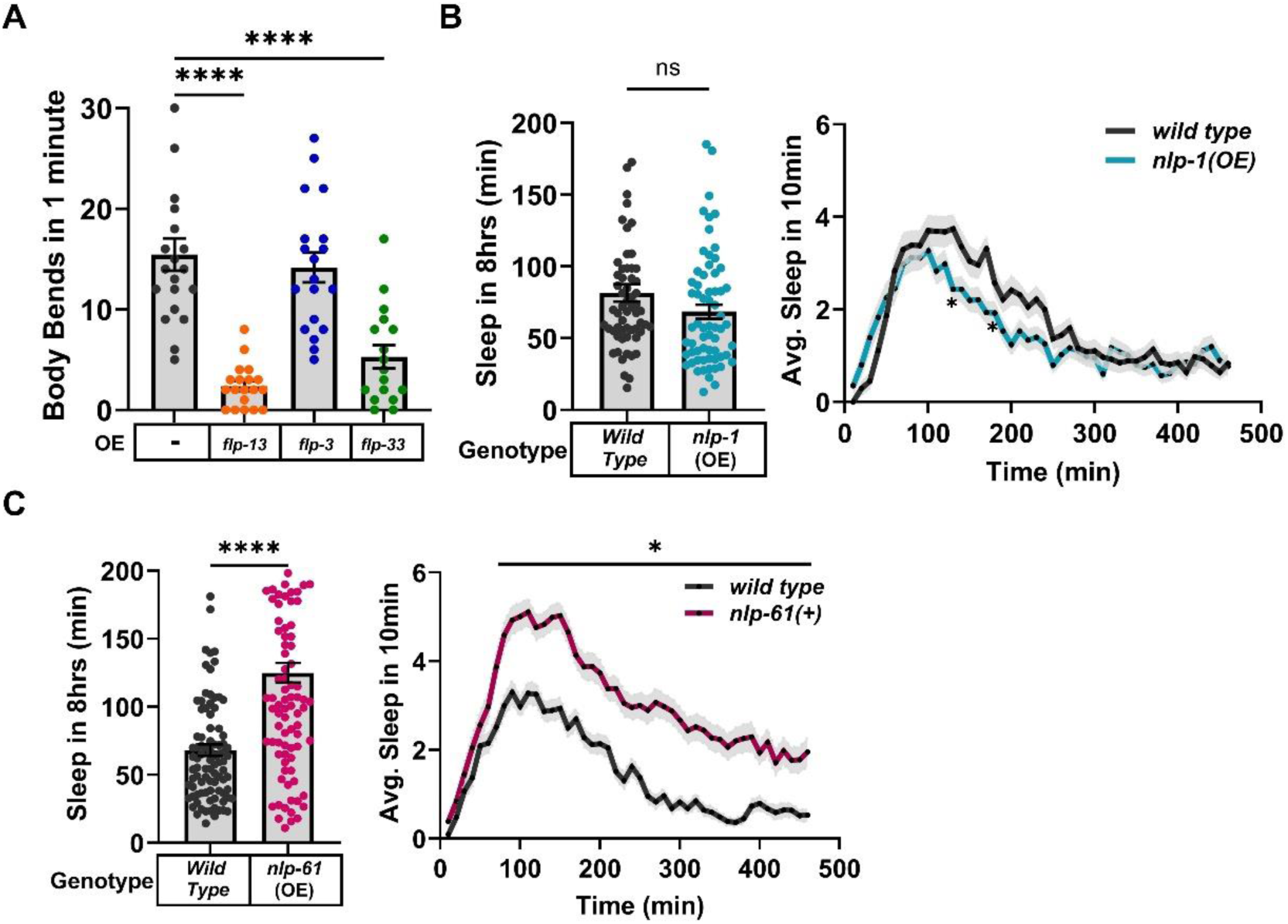
Over expression of some neuropeptides can alter sleep. **A)** Average number of body bends in one minute, two-hours post heat shock (30 minutes at 33°C), in wild-type (N=20), and transgenic strains over-expressing *flp-13* (N=19), *flp-3* (N=17), and *flp-33* (N=17). Statistical analyses were performed by one-way ANOVA followed by Tukey’s test (****p<0.0001). **B)** Quiescence in wild-type (N=60) and *nlp-1*(OE) (N=64) animals. **C)** Quiescence in wild-type (N=91) and *nlp-61*(OE) (N=94) animals. Right panel – *p<0.05 at 70-390, 410, 430, and 450-460 minutes. For **B) and C)** Student’s t-test was used to compare total quiescence (Left panels) and two-way ANOVA followed by Sidak’s test was used for temporal analyses (Right panels) (*p<0.05, ****p<0.0001).

**Figure S2:**
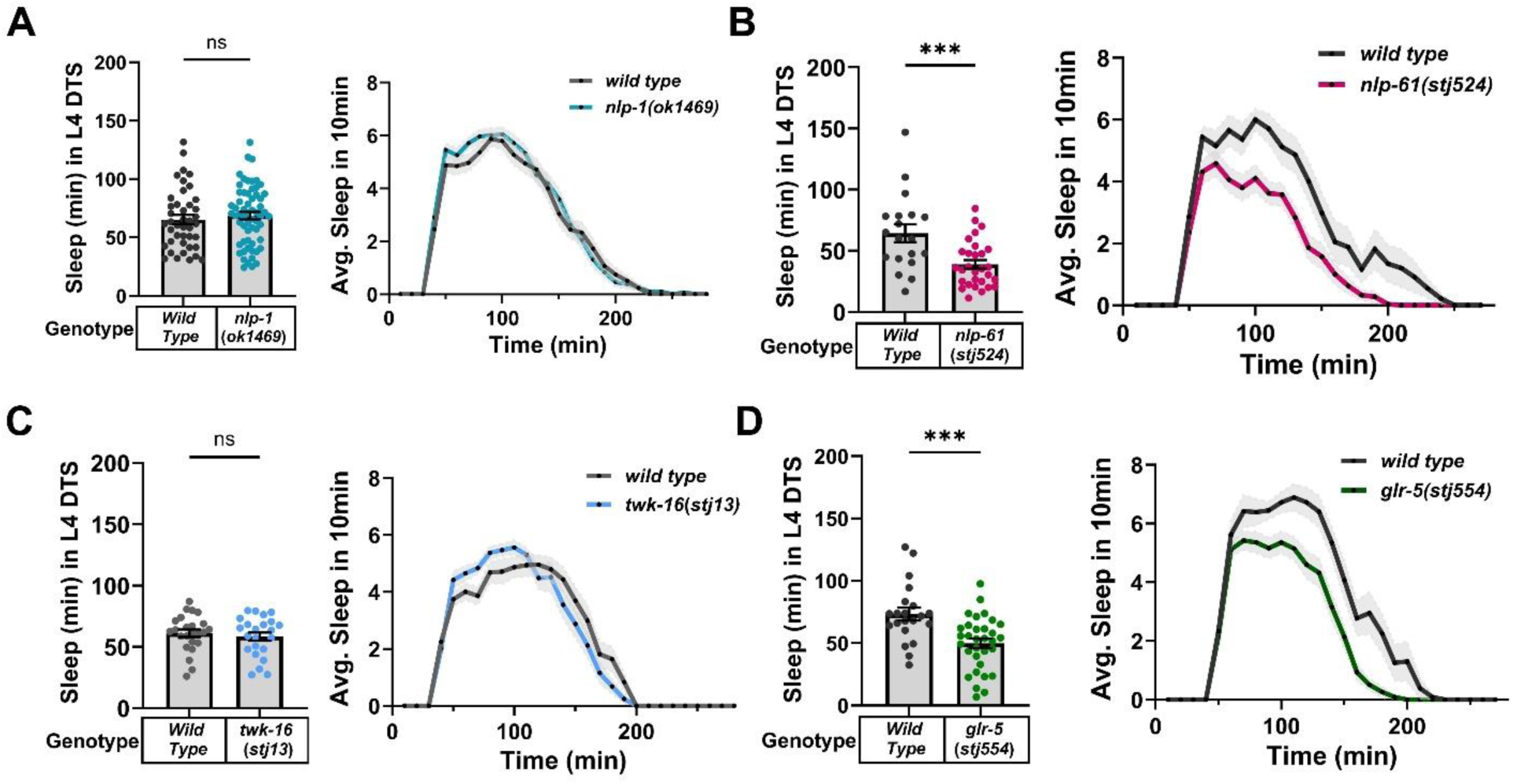
Developmentally-timed sleep of *nlp-1*, *nlp-61*, *twk-16*, and *glr-5* mutants. For each panel, the average total minutes of movement quiescence during L4 DTS is displayed on the left, and the average minutes of movement quiescence in 10-minute windows during L4 DTS on the right. **A)** Quiescence in wild-type (N=42) and *nlp-1*(*ok1469*) (N=61) animals. **B)** Quiescence in wild-type (N=19) and *nlp-61*(*stj524*) (N=29) animals. **C)** Quiescence in wild-type (N=23) and *twk-16*(*stj13*) (N=24) animals. **D)** Quiescence in wild-type (N=21) and *glr-5*(*stj554*) (N=34) animals. Student’s t-test was used for comparison of total sleep between genotypes, and temporal comparisons were made using two-way ANOVA and Sidak’s test (***p<0.001).

**Figure S3:**
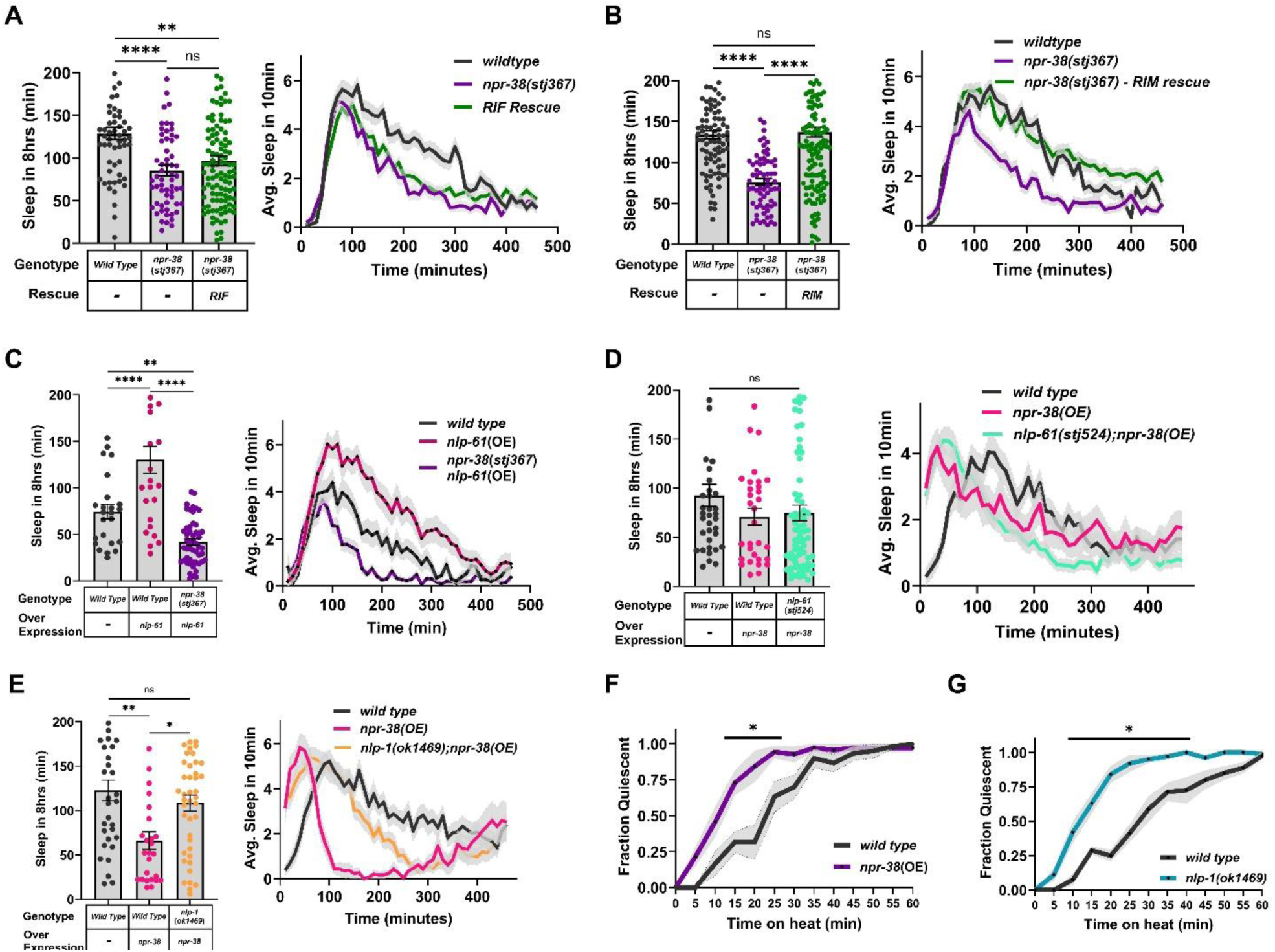
Other timing defects related to *nlp-1* and *npr-38*. **A)** Quiescence in wild-type (N=58), *npr-38*(*stj367*) (N=60), and *npr-38*(*stj367*) transgenic (RIF rescue; Promoter: *nlp-6*) (N=110) animals. **B)** Quiescence in wild-type (N=92), *npr-38*(*stj367*) (N=66), and *npr-38*(*stj367*) transgenic (RIM rescue; Promoter: *cex-1*) (N=145) animals. **C)** Quiescence in wild-type (N=24), *nlp-61*(OE) (N=24), and *npr-38*(*stj367*) transgenic (*nlp-61*(OE)) (N=51) animals. **D)** Quiescence in wild-type (N=37), *npr-38*(OE) (N=31), and *nlp-61*(*stj524*) transgenic (*npr-38*(OE)) (N=70) animals. **E)** Quiescence in wild-type (N=33), *npr-38*(OE) (N=26), and *nlp-1*(*ok1469*) transgenic (*npr-38*(OE)) (N=42) animals. For **A-E**, statistical significance was calculated by one-way ANOVA followed by Tukey’s test (*p<0.05, **p<0.01, ***p<0.001, ****p<0.0001). **F)** Average fraction of quiescent animals in the presence of noxious heat (37°C) in wild-type (6 trials, N=10 animals each) and *npr-38*(OE) animals (7 trials, N=10 animals each). **G)** Average fraction of quiescent animals in the presence of noxious heat (37°C) in wild-type (8 trials, N=10 animals each) and *nlp-1*(*ok1469*) animals (10 trials, N=10 animals each). Significance was calculated by two-way ANOVA followed by Sidak’s multiple comparisons test (*p<0.05).

**Figure S4:**
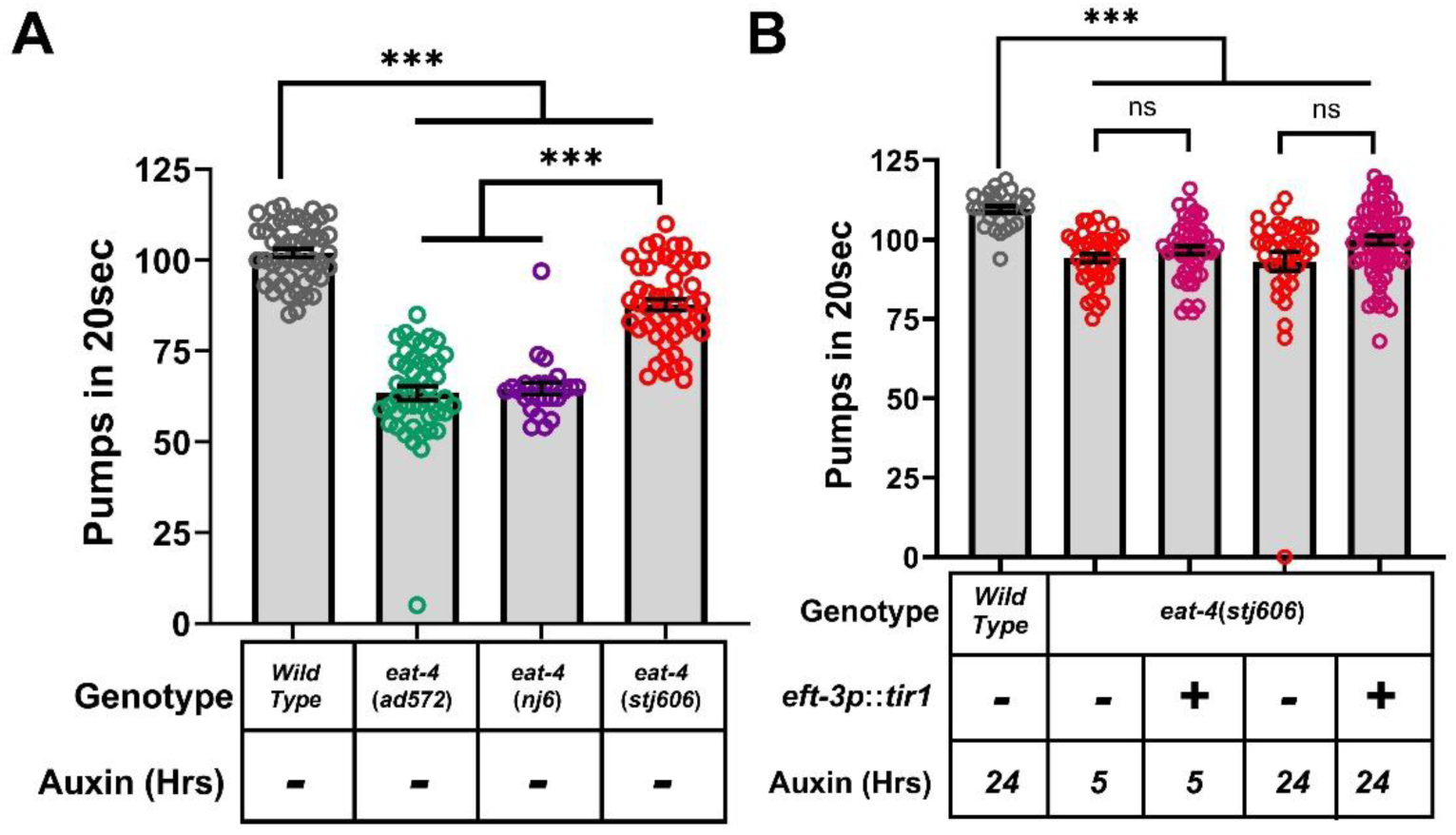
Pumping rate of *eat-4* mutants. **A)** Average pumps in 20 seconds in wild-type (N=48), *eat-4*(*ad450*) (N=56), *eat-4*(*nj6*) (N=51), and *eat-4*(*stj606*) (N=50) animals in the absence of auxin. **B)** Average pumps in 20 seconds in wild-type, *eat-4*(*stj606*), and transgenic *eat-4*(*stj606*) animals expressing *tir1* in all cells driven from the *eft-3* promoter (*eft-3p*::*tir1*), in the presence of auxin for the time-period displayed below the bars. One-way ANOVA followed by Tukey’s test was used to make comparisons between genotypes and conditions (***p<0.001).

**Figure S5:**
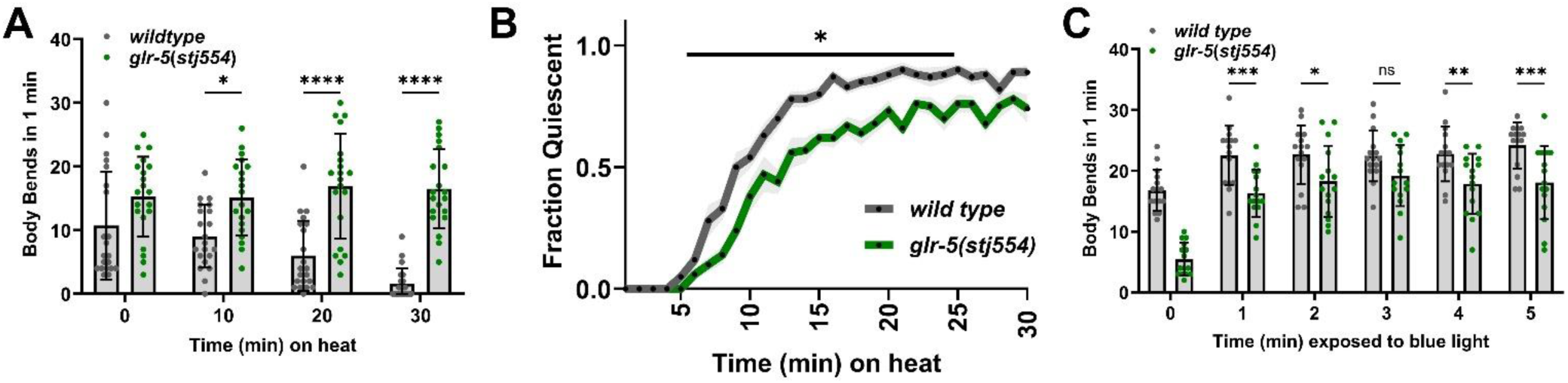
The response to acute stress in *glr-5*(*stj554*) mutants. **A)** Average body bends performed in 1 minute following exposure to a 37°C heat stress for the designated number of minutes (min). **B)** Average fraction of quiescent animals in the presence of noxious heat (37°C) in wild-type (10 trials, N=10 animals each) and *glr-5*(*stj554*) animals (10 trials, N=10 animals each). Significance was calculated by two-way ANOVA followed by Sidak’s multiple comparisons test (*p<0.05). **C)** Average body bends performed in 1 minute (min) in the presence of blue light. For **A** and **C**, Student’s t-test was used to make comparisons between genotype (*p<0.05, **p<0.01,***p<0.001, ****p<0.0001). For **B**, significance was calculated by two-way ANOVA followed by Sidak’s multiple comparisons test (*p<0.05).

**Figure S6:**
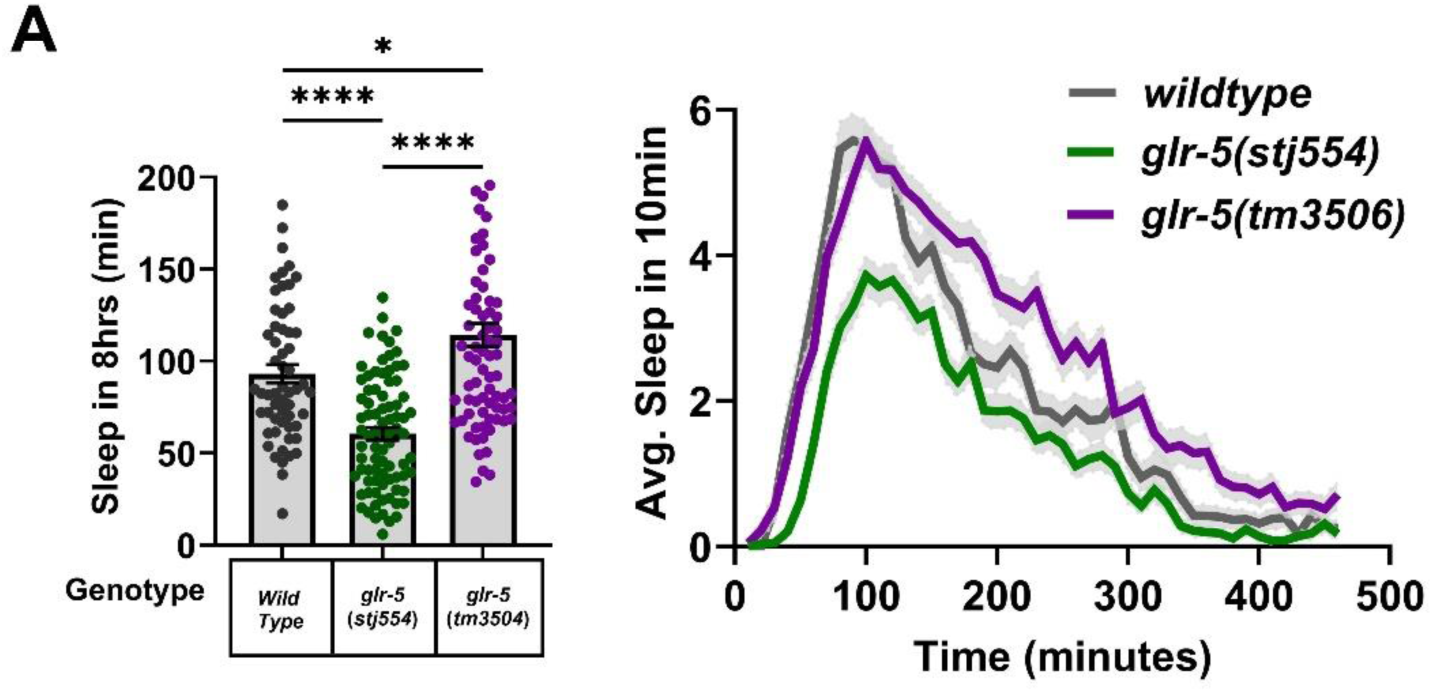
*glr-5*(*tm3506*) mutants display enhanced sleep. **A)** Quiescence in wild-type (N=52), *glr-5*(*stj554*) (N=77), and *glr-5*(*tm3506*) (N=87) animals. Statistical significance was calculated by one-way ANOVA followed by Tukey’s test (*p<0.05, ****p<0.0001).

**Figure S7:**
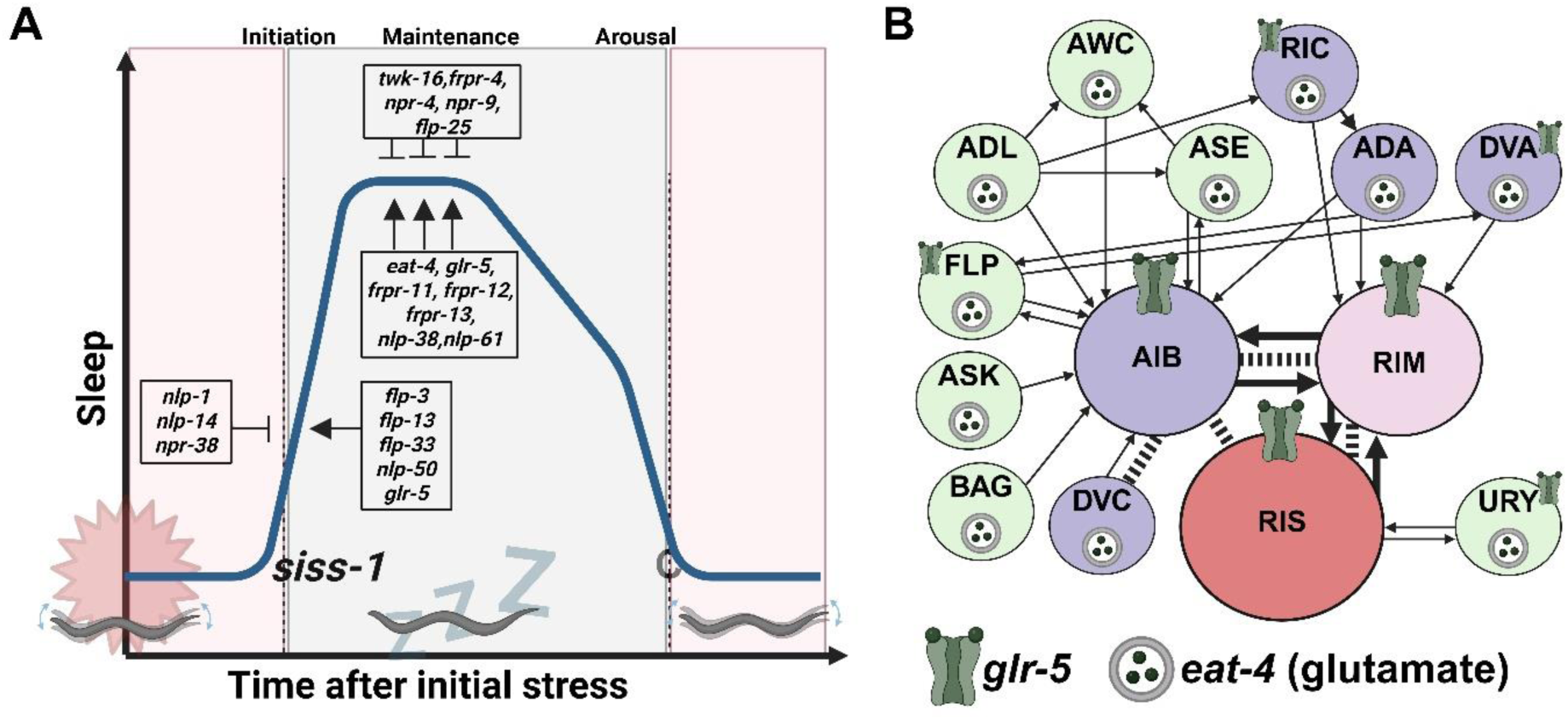
A proposed neuropeptidergic and glutamatergic model of sleep regulation. **A)** A schematic of a proposed role for each gene during sleep initiation and/or maintenance. **B)** Glutamatergic neurons (*eat-4*-expressing) with direct synaptic connections to AIB, RIM, and RIS.

**Table S1:** Strains used in the current study.

| Strain Name | Genotype | Description | Location |
| --- | --- | --- | --- |
| VC2497 | *flp-3(ok3265)* | 457bp deletion | CGC |
| NQ602 | *flp-13(tm2427)* | 380bp deletion | Raizen Lab |
| VC1982 | *flp-25(gk1016)* | 673bp deletion | CGC |
| VC2422 | *flp-33(gk1037)* | 507bp deletion | CGC |
| SJU322 | *flp-3(ok3265); flp-33(gk1037)* | VC2397 crossed with VC2422 | Nelson Lab |
| RB1340 | *nlp-1(ok1469)* | 700bp deletion | CGC |
| RB1341 | *nlp-1(ok1470)* | 600bp deletion | CGC |
| VC2357 | *nlp-38(ok2330)* | 1000bp deletion | CGC |
| TM1782 | *npr-4(tm1782)* | 1226bp deletion | NBRP |
| VC4220 | *frpr-16(gk5305[loxP;myo-2p::gfp::unc-54 3' UTR; rps-27p::neoR::unc-54 3' UTR; loxP])* | 1685bp deletion | CGC |
| TM1652 | *npr-9(tm1652)* | 318 bp deletion + 2 bp insertion | NBRP |
| RB1365 | *npr-16(ok1541)* | 145bp deletion | CGC |
| CHS1058 | *npr-12(yum1406); npr-9(yum1404); npr-11(yum1405)* | Frameshift deletion and insertion of stop codon (Pu *et al.* 2023) | CGC |
| IK602 | *eat-4(nj6)* | Substitution – loss of function | CGC |
| DA572 | *eat-4(ad572)* | Substitution – loss of function | CGC |
| TM3506 | *glr-5(tm3506)* | 245bp deletion | NBRP |
| NQ385 | *frpr-4(ok2376)* | 1540bp deletion | Raizen Lab |
| NQ769 | *frpr-6(tm6135)* | 889 bp deletion | Nelson Lab |
| NQ915 | *dmsr-1(qn45)* | 278bp indel | Raizen Lab |
| SJU210 | *twk-16(stj13)* | 7bp insertion in exon 4 – tgctagc | Nelson Lab |
| SJU253 | *frpr-4(stj21)* | 6bp insertion | Nelson Lab |
| SJU261 | *twk-16(stj16)* | 7bp insertion in exon 6 – tgctagc | Nelson Lab |
| SJU269 | *twk-16(stj13); stjEx181[twk-16(+); myo-2p::mCherry]* | Rescue of *twk-16* with a genomic PCR fragment spanning ~8kb (5'UTR, coding, and 3'UTR) | Nelson Lab |
| SJU277 | *frpr-11(stj22); frpr-12(stj23)* | *frpr-11* – 6bp insertion in exon 4 – gctagc<br>*frpr-12* – 7bp insertion in exon 4 – agctagc | Nelson Lab |
| SJU285 | *stjEx185[hsp-16.2p::flp-3; myo-2p::mCherry]* | *flp-3* overexpression | Nelson Lab |
| SJU287 | *stjEx187[hsp-16.2p::flp-33; myo-2p::mCherry]* | *flp-33* overexpression | Nelson Lab |
| SJU361 | *stjEx227[npr-38(+); myo-2p::gfp]* | *npr-38* overexpression |  |
| SJU471 | *frpr-11(stj22); frpr-12(stj23); frpr-13(stj458)* | *frpr-13* – 9bp insertion in exon 3 – gctagctag | Nelson Lab |
| SJU524 | *nlp-61(stj524)* | 12bp insertion in exon 1 – taagctagctaa | Nelson Lab |
| SJU554 | *glr-5(stj554)* | 13bp insertion in exon 3 – taattaggctagc | Nelson Lab |
| SJU561 | *glr-5(stj554); stj351[flp-21p::glr-5; myo-2p::mCherry]* | RMG rescue – *glr-5* | Nelson Lab |
| SJU562 | *glr-5(stj554); stj352[flp-21p::glr-5; myo-2p::mCherry]* | RMG rescue – *glr-5* | Nelson Lab |
| SJU573 | *stjEx362[nlp-61(+); myo-2p::mCherry]* | *nlp-61* overexpression | Nelson Lab |
| SJU578 | *glr-5(stj554); stjEx366[ver-3p::glr-5; flp-11p::glr-5; myo-2p::mCherry]* | ALA and RIS rescue – *glr-5* | Nelson Lab |
| SJU581 | *glr-5(stj554); stjEx369[glr-5(+); myo-2p::mCherry]* | Genomic rescue – *glr-5* | Nelson Lab |
| SJU594 | *glr-5(stj554); stjEx379[npr-9p::glr-5; myo-2p::mCherry]* | AIB rescue – *glr-5* | Nelson Lab |
| SJU595 | *glr-5(stj554); stjEx380[npr-9p::glr-5; myo-2p::mCherry]* | AIB rescue – *glr-5* | Nelson Lab |
| SJU606 | *eat-4(stj606)* | Insertion of degron sequence (ZHANG *et al.* 2015) in place of stop codon at 3' end | Nelson Lab |
| SJU611 | *nlp-61(stj524); stjEx227[npr-38(+); myo-2p::gfp]* | *nlp-61* *lof* and *npr-38* overexpression | Nelson Lab |
| SJU614 | *nlp-1(ok1469); stjEx227[npr-38(+); myo-2p::gfp]* | *nlp-1* *lof* and *npr-38* overexpression | Nelson Lab |
| SJU627 | *stjEx404[nlp-1(+); myo-2p::mCherry]* | *nlp-1* overexpression | Nelson Lab |
| SJU638 | *npr-38(stj367); stjEx412[cex-1p::npr-38; myo-2p::mCherry]* | RIM rescue – *npr-38* | Nelson Lab |
| SJU639 | *npr-38(stj367); stjEx413[cex-1p::npr-38; myo-2p::mCherry]* | RIM rescue – *npr-38* | Nelson Lab |
| SJU641 | *glr-5(stj554); stjEx415[twk-16p::glr-5; myo2p::mCherry]* | DVA rescue – *glr-5* | Nelson Lab |
| SJU644 | *glr-5(stj554); stjEx418[cex-1p::glr-5; myo-2p::mCherry]* | RIM rescue – *glr-5* | Nelson Lab |
| SJU651 | *npr-38(stj367); stjEx425[nlp-6p::npr-38; myo-2p::mCherry]* | RIF rescue – *npr-38* | Nelson Lab |
| SJU660 | *glr-5(stj554); stjEx429[aptf-1p::glr-5; cex-1p::glr-5; myo-2p::mCherry]* | AIB, RIS, and RIM rescue – *glr-5* | Nelson Lab |

**Table S2:**
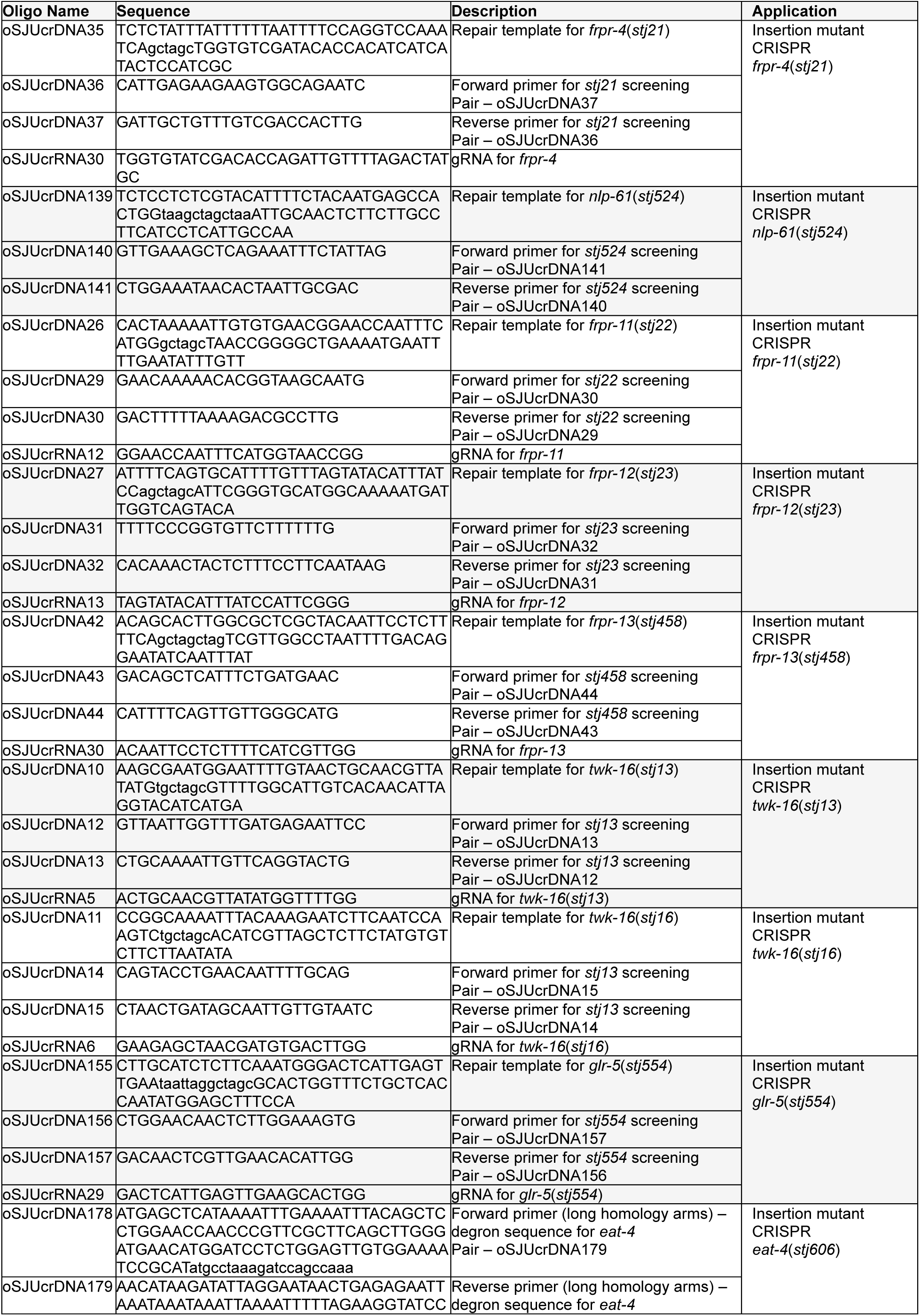
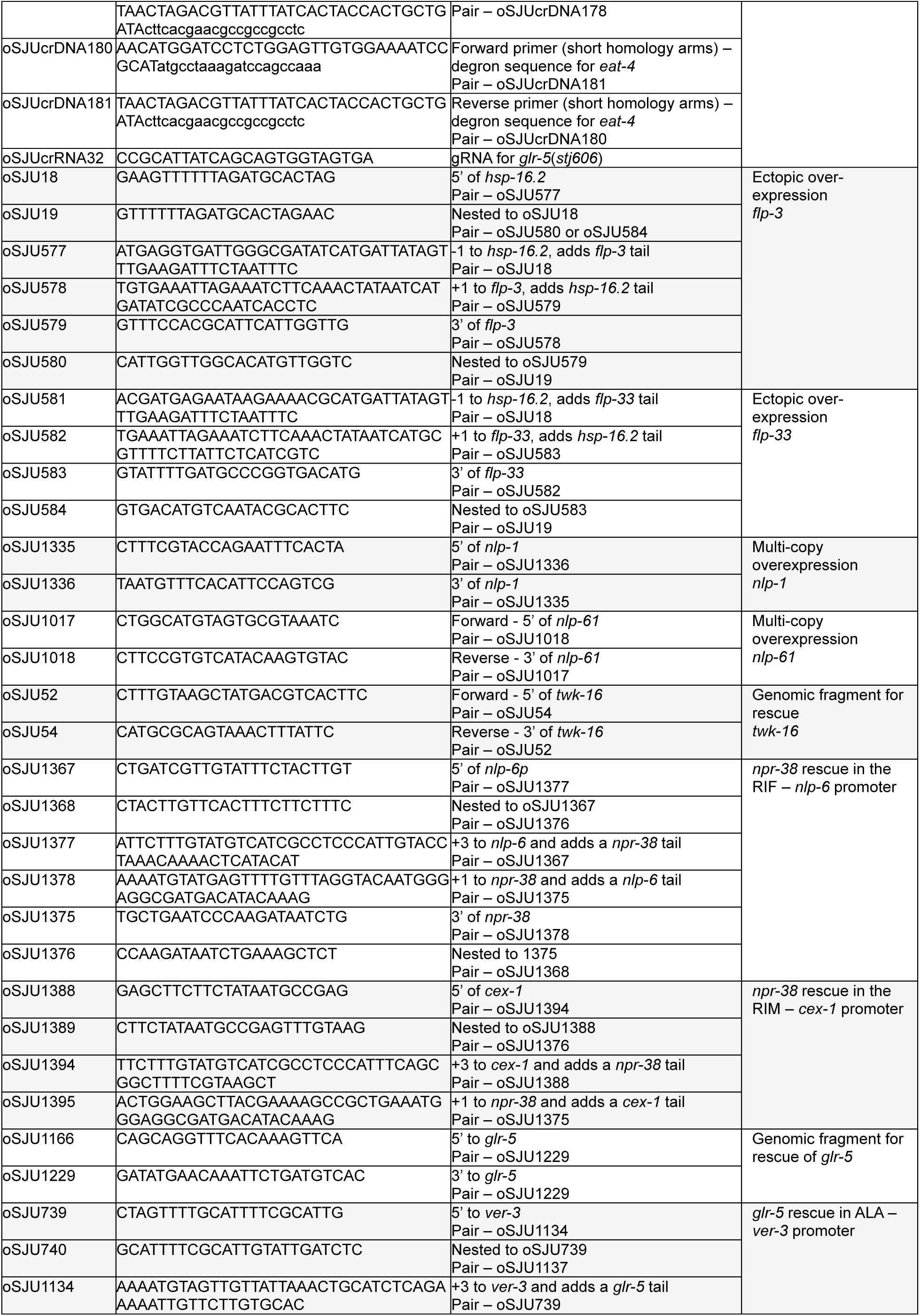
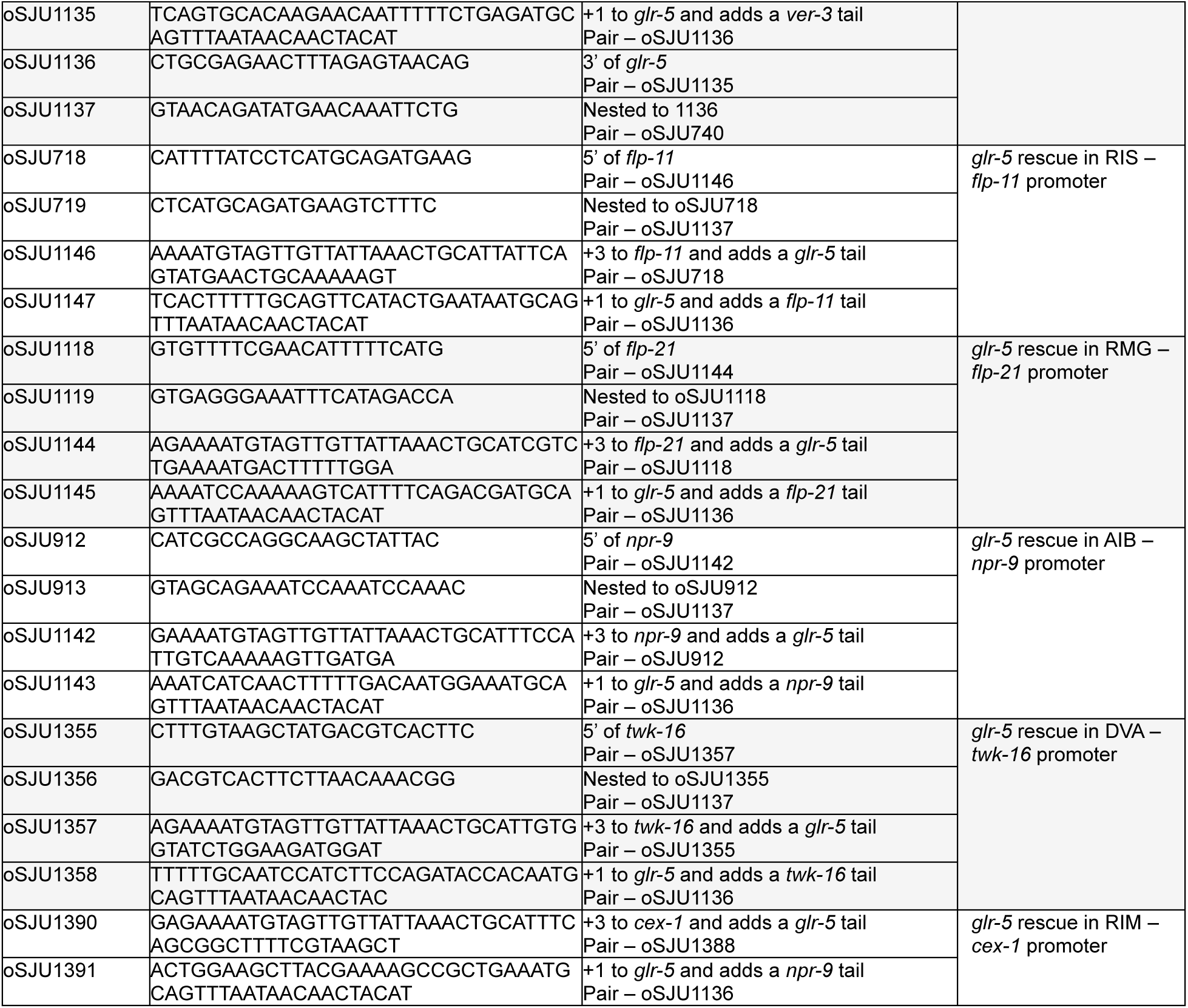
Oligonucleotides used in the current study (Integrated DNA Technologies).

